# Spatiotemporal map of the developing human reproductive tract at single-cell resolution

**DOI:** 10.1101/2024.10.30.621114

**Authors:** Valentina Lorenzi, Cecilia Icoresi Mazzeo, Nadav Yayon, Elias R. Ruiz-Morales, Carmen Sancho-Serra, Frederick C.K. Wong, Magda Marečková, Liz Tuck, Kenny Roberts, Tong Li, Marc-Antoine Jacques, Xiaoling He, Roger Barker, Berta Crespo, Batuhan Cakir, Simon Murray, Martin Prete, Yong Gu, Iva Kelava, Luz Garcia Alonso, John C Marioni, Roser Vento Tormo

**Author notes:** equal contribution (Valentina Lorenzi, Cecilia Icoresi Mazzeo).

## Abstract

The human reproductive tract plays an essential role in species perpetuation. Its development involves complex processes of sex specification, tissue patterning and morphogenesis, which, if disrupted, can cause lifelong health issues, including infertility. Here, we generated an extensive single-cell and spatial multi-omic atlas of the human reproductive tract during prenatal development, which allowed us to answer questions that smaller-scale, organ-focused experiments could not address before. We identified potential regulators of sexual dimorphism in reproductive organs, pinpointing novel genes involved in urethral canalisation of the penis, with relevance to hypospadias. By combining histological features with gene expression data, we defined the transcription factors and cell signalling events required for the regionalisation of the Müllerian and Wolffian ducts. This led to a refinement of how the *HOX* code is established in the distinct reproductive organs, including increased expression of thoracic *HOX* genes in the rostral mesenchyme of the fallopian tube and epididymis. Our study further revealed that the epithelial regionalisation of the fallopian tube and epididymis required for sperm maturation in adulthood is established early in development. In contrast, later events in gestation or postnatally are necessary for the regionalisation of the uterocervical canal epithelium. By mapping sex-specific reproductive tract regionalisation and differentiation at the cellular level, our study offers valuable insights into the causes and potential treatments of reproductive disorders.

## Introduction

The development of the human reproductive tract is a complex morphogenetic process orchestrated by paracrine interactions^1,2^ and hormonal signalling^3,4^. The internal genitalia (with the exception of the gonads) originate from the Müllerian and Wolffian ducts and the urogenital sinus. In females (XX), the Müllerian ducts develop into the fallopian tubes, uterus, cervix and upper vagina, while the urogenital sinus forms the lower vagina^5,6^. In males (XY), the Wolffian ducts give rise to the epididymis, vas deferens and seminal vesicles, while the urogenital sinus becomes the prostate^7,8^. The genital tubercle gives rise to the external genitalia: the clitoris in females and the penis in males^9^.

Initially, embryonic reproductive tissue precursors (i.e. Müllerian/Wolffian ducts, urogenital sinus and genital tubercle) comprise an undifferentiated epithelial inner layer and surrounding mesenchyme. As development progresses, the sexually dimorphic differentiation of the mesenchyme precedes and dictates the differentiation of the epithelium^1^. Müllerian and Wolffian duct differentiation is particularly complex, as precise spatial boundaries must be established between the resulting organs^10,11^.

Both Müllerian and Wolffian ducts are present in genetically female and male embryos until approximately 9-10 post-conceptional weeks (PCW). If the embryonic gonads differentiate into testes under the control of the Y chromosome-linked *SRY* gene^12,13^, Sertoli cells in the testes produce anti-Müllerian hormone (AMH), which causes the Müllerian ducts to regress^14–17^. Leydig cells in the testes secrete testosterone, which promotes the development of the Wolffian ducts into the upper reproductive tract^7^, and which is further converted into dihydrotestosterone, masculinising the lower reproductive tract^9,18^. In the absence of *SRY* and these hormones, as in XX embryos, the Wolffian ducts regress and the Müllerian ducts, urogenital sinus and genital tubercle develop into the female reproductive tract.

Genetic and environmental disruptions to reproductive tract development can lead to congenital anomalies, infertility and cancer^19,20^. For example, approximately 7% of women have congenital uterine anomalies, rising to ∼17% among those with recurrent miscarriages^19^. However, the cellular and molecular mechanisms mediating human reproductive tract development remain poorly studied and have primarily been inferred from rodent and chicken loss-of-function studies or human histological observations^21–26^. Recently, we and others have used single-cell transcriptomics to study the developing human reproductive tract, primarily focussing the gonads^27–30^ and, to a limited extent, the upper portions of the Müllerian and Wolffian ducts^31^. However, we lack a holistic study of the entire tract development in both sexes.

Here, we generated a highly-resolved, spatiotemporal, multi-omic map of the entire developing human reproductive tract (excluding the gonads), profiling over half a million cells spanning the first and second trimesters. We detailed the cellular and molecular features of the female and male reproductive tracts throughout the critical stages of sexual differentiation, revealing how sex-specific signals drive the dimorphic development of reproductive organs and the regression of sexually unmatched ducts. Additionally, we resolved the cascade of gene expression changes from tissue-wide gradients into lineage-specific compartments within the Müllerian and Wolffian ducts, defining the key transcription factors and cell-cell communication events that drive their differentiation into final organ derivatives. Finally, we harness our atlas to pinpoint cell types and developmental windows likely affected by endocrine-disrupting chemicals and clinically approved drugs.

## Results

### Spatiotemporal atlas of human reproductive tract development

We profiled 80 reproductive tract samples from 6 to 21 PCW, covering stages of specification and differentiation of the internal and external genitalia as well as the regression of unmatched reproductive ducts. Our study employed single-cell RNA sequencing (scRNA-seq, 471,749 cells), single-cell chromatin accessibility sequencing (scATAC-seq, 175,227 cells), spatially resolved gene expression profiling via *In Situ* Sequencing (*ISS*, 11 slides) and *10x Visium* (25 slides) (**Figure 1a-b**). By mapping the dissociated single-cell data onto stage-matched *ISS* and *10x Visium* data, we enhanced the resolution and stringency of cell type definitions, identifying 52 distinct reproductive tract-specific cell types (**Figure 1c**). The integration of spatially-resolved data was crucial for cell annotation as unique markers for many of the identified cell types had not been previously described (**Methods**, **Supplementary Note 1**).

**Figure 1.**
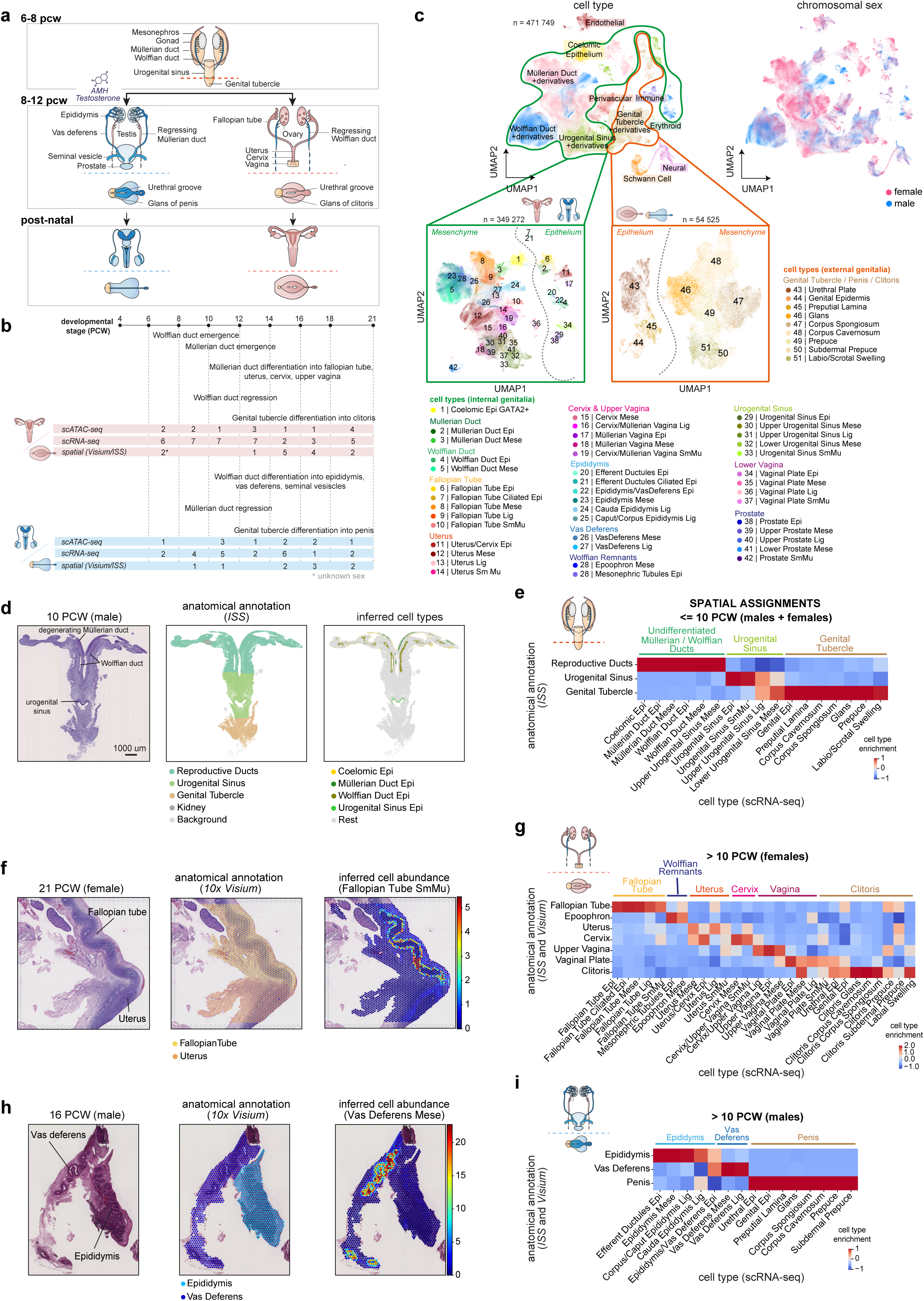
Spatiotemporal atlas of human reproductive tract development. **a,** Schematic illustration of human reproductive development showing the main anatomical structures in XX and XY embryos and fetuses. **b,** Diagram summarising the stage and sex composition of our sample cohort along with main events occurring during reproductive tract development. **c,** (top left) Batch-corrected Uniform Manifold Approximation and Projection (UMAP) embedding of the scRNA-seq dataset (n = 471,749 cells) coloured by major developmental lineage and (top right) chromosomal sex. (bottom left) Batch-corrected UMAP embedding of the reproductive-specific scRNA-seq cells from the internal genitalia (n = 349,272 cells) coloured by cell type. (bottom right) UMAP embedding of the scRNA-seq cells from the external genitalia (n = 54,525 cells) coloured by cell type. **d,** (left) Hematoxylin and eosin (H&E) stained image of a representative 10 PCW male sample profiled with *In Situ* Sequencing (*ISS*). (middle) Annotations of the anatomical structures using *TissueTag*^60^ and (right) inferred cell type labels for selected cell types from the scRNA-seq dataset in the *ISS* slide using *iss-patcher*^103^. **e,** Heatmap showing the z-score enrichment of each epithelial, mesenchymal, smooth muscle and ligament cell type annotated in early male and female scRNA-seq samples <=10 PCW (x-axis) in *ISS* cells corresponding to each anatomical annotation (y-axis). **f,** (left) H&E stained image of a representative 21 PCW female sample profiled with *10x Visium*. (middle) Annotations of the anatomical structures using *TissueTag*^60^ and (right) spatial mapping of fallopian tube smooth muscle cells from the scRNA-seq dataset onto the corresponding *10x Visium* slide using *cell2location*^104^. Estimated cell type abundance (colour intensity) in each *10x Visium* spot is shown over the H&E image. **g,** Heatmap showing the combined z-score enrichment of each epithelial, mesenchymal, smooth muscle and ligament cell type annotated in female scRNA-seq samples >10 PCW (x-axis) in *10x Visium* spots and *ISS* cells corresponding to each anatomical annotation (y-axis). **h,** (left) H&E stained image of a representative 16 PCW male sample profiled with *10x Visium*. (middle) Annotations of the anatomical structures using *TissueTag*^60^ and (right) spatial mapping of vas deferens mesenchymal cells from the scRNA-seq dataset onto the corresponding *10x Visium* slide using *cell2location*^104^. Estimated cell type abundance (colour intensity) in each *10x Visium* spot is shown over the H&E image. **i,** Heatmap showing the combined z-score enrichment of each epithelial, mesenchymal, smooth muscle and ligament cell type annotated in male scRNA-seq samples >10 PCW (x-axis) in *10x Visium* spots and *ISS* cells corresponding to each anatomical annotation (y-axis).

In the early stages of development (until ∼9/10 PCW), we identified Müllerian (*WNT7A*/*SOX17+*^32,33^ epithelium, *CNTN1+* mesenchyme) and Wolffian (*WNT9B*/*GATA3+*^34,35^ epithelium, *PLAC1/HTR2B+* mesenchyme) duct cells (**Figure 1 c-e**, **Supplementary Figure 1 a-c, Supplementary Note 1**). Additionally, we detected cells from the urogenital sinus (*FOXA1*/*SHH+* epithelium, *RDH10+* mesenchyme) and genital tubercle (*UPK1A/B+* epithelium, *TBX4/5+* mesenchyme) (**Figure 1 c-e**, **Supplementary Figure 1 a-c, Supplementary Note 1**). In samples from <8 PCW embryos, various cell types from adjacent kidney and adrenal glands are also present, as complete tissue microdissection at this developmental stage is challenging (**Supplementary Figure 1d-e**).

As gestation progresses (9-21 PCW), female-specific cells emerge in the fallopian tubes (*PNOC/ERP27+* non-ciliated and *DNAH12+* ciliated epithelium, *LGR5/TSPAN8+* mesenchyme), uterocervix (*UCA1/DLX5+* epithelium, *ITGA4/RORB+* mesenchyme), and vagina (*DLX5/TP63+* Müllerian-derived epithelium, *FOXA1/PRAC1+* urogenital sinus-derived epithelium, *SRD5A2+* mesenchyme) in XX samples (**Figure 1 c, f-g**, **Supplementary Figure 2a-f, Supplementary Note 1**). Notably, the uterus and cervix show a very similar cell type composition, indicating that further regionalisation likely occurs after 21 PCW (**Figure 1g**). In the same time window (9-21 PCW), male-specific epididymis (*PDZK1/GLYAT+* non-ciliated and *DNAH12+* ciliated epithelium, *PLAC1/HTR2B+* mesenchyme) and vas deferens (*MUC6/MARCH11*+ epithelium, *RAI2/CHD7+* mesenchyme) cells appear in XY samples (**Figure 1c, h-i**, **Supplementary Figure 3a-d, Supplementary Note 1**).

Our atlas also captures the remnants of the sexually unmatched reproductive ducts that persist in both sexes. Wolffian-like mesenchymal and epithelial cells (epoophoron) are apparent near the fallopian tubes in females until 21 PCW, while fallopian-like epithelial cells are observed in some male samples between 10 and 14 PCW, indicating that Müllerian differentiation can occur in males before regression orchestrated by AMH concludes (**Figure 1c, f-g**, **Supplementary Figure 2a-b, e**, **Supplementary Figure 3a-b,e**).

Consistent with studies in the mouse^36,37^, we do not identify sex-specific cell types in the developing penis and clitoris. Across all developmental stages and in both sexes we identify the urethral epithelium (*FOXA1/SHH+*), erectile tissues (*corpus cavernosum* (*SOX9/PRR16+*) and *corpus spongiosum* (*FOXF1/SALL1+*)), glans (*SP9/MSX1+*), prepuce (*SIX1/SHOX2+*) and genital epidermis (*KRT14/KRTDAP+*) (**Supplementary Figure 4a,c, Supplementary Note 1**). Moreover, a mixture of endodermal (*FOXA1*) and ectodermal (*KRT14*) markers is observed at the urethral meatus (**Supplementary Figure 4b,d**).

In summary, our spatio-temporal, single-cell resource represents the most comprehensive and unbiased characterisation of the reproductive epithelia and surrounding mesenchyme during human prenatal development, covering the progression from undifferentiated precursors to differentiated sex-specific organs (accessible at www.reproductivecellatlas.org).

### Sexual dimorphism in the genital tubercle

We next investigated how sexual dimorphism is acquired in individual reproductive tract organs, beginning with the external genitalia. Androgen action is known to lead to an increase in size of the penis and canalisation of the male urethra^9,25,38,39^ (**Figure 2a**). The mesenchymal *corpus spongiosum*, adjacent to the invaginating urethral epithelium, plays a crucial role in urethral canalisation by moving medially and shaping the urethral canal in the developing penis^40^.

**Figure 2.**
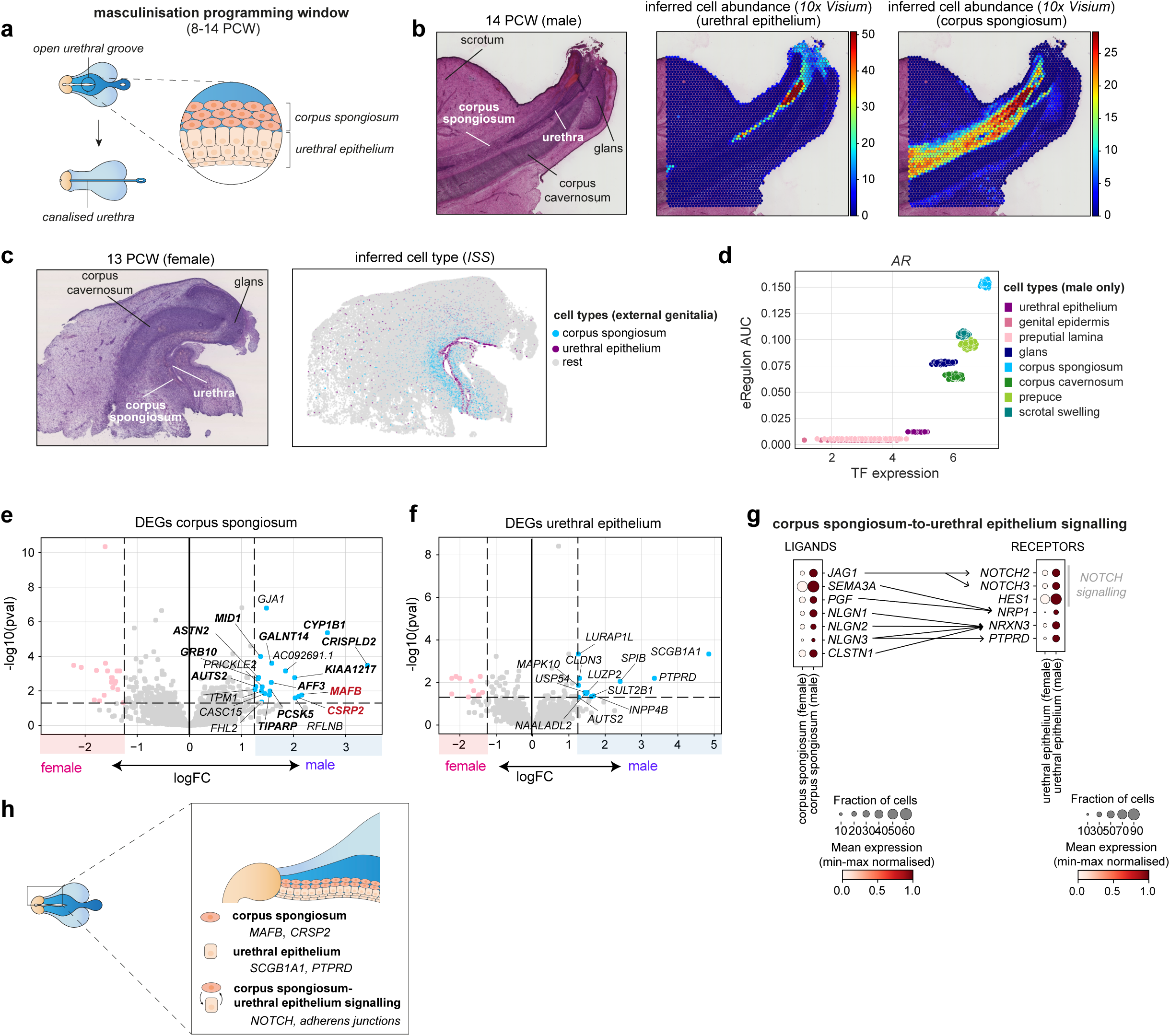
Sexual dimorphism in the genital tubercle. **a,** Schematic illustration of the process of urethral canalisation that occurs in the male genital tubercle during the masculinisation programming window (∼8-14 PCW). **b,** (left) Hematoxylin and eosin (H&E) stained image of a representative 14 PCW male sample profiled with *10x Visium* alongside (right) spatial mapping of urethral epithelium and *corpus spongiosum* cells from the scRNA-seq dataset onto the corresponding *10x Visium* slide using *cell2location*^104^. Estimated cell type abundance (colour intensity) in each *10x Visium* spot is shown over the H&E image. **c,** (left) H&E stained image of a representative 13 PCW female sample profiled with *In Situ* Sequencing (*ISS)* alongside (right) inferred cell type labels for urethral epithelium and *corpus spongiosum* from the scRNA-seq dataset in the corresponding *ISS* slide using a *iss-patcher*^103^. **d,** Scatter plot showing the area under the curve of the inferred activity of the chromatin region-based cistrome for *AR* (y-axis) as a function of its expression (x-axis) in each scATAC/RNA-seq metacell derived from the male genital tubercle during the masculinisation programming window (8-14 PCW). Each metacell is coloured by cell type. **e,** Volcano plot showing the log fold-change (x-axis) and adjusted p-value (y-axis) of the differential expression test (*pyDESEQ2*^105^: adjusted p-value = 0.05, |logFC| > 1.25) between male and female samples within the human *corpus spongiosum*. Only upregulated genes in males are labelled in the plot. **f,** Volcano plot showing the log fold-change (x-axis) and adjusted p-value (y-axis) of the differential expression test (*pyDESEQ2*^105^: adjusted p-value = 0.05, |logFC| > 1.25) between male and female samples within the human urethral epithelium. Only upregulated genes in males are labelled in the plot. **g,** Dotplot showing the log-transformed, min-max normalised expression of *corpus spongiosum* ligands and urethral epithelium receptors (y-axis) in each cell type (x-axis), separated by sex. Interacting ligands and receptors are connected by an arrow. **h,** Schematic illustration summarising the putative drivers of urethral canalisation found by our analysis.

Focusing on samples from 8 to 14 PCW, corresponding to the masculinisation programming window (MPW) — the critical period during which disruptions in androgen signalling have the most significant phenotypic effects on newborn males^41^ — we first validated the identity of the urethral epithelium and surrounding *corpus spongiosum* in both the penis and clitoris through spatial mapping (**Figure 2b,c**). In the developing penis, the *corpus spongiosum* showed the highest androgen receptor (*AR*) activity (as evaluated by expression and chromatin accessibility) compared to all other cell types (**Figure 2d, Supplementary Figure 5a-b**).

To identify candidate target genes downstream of androgen signalling involved in urethral canalisation, we performed differential expression analysis within the mesenchymal *corpus spongiosum* between male and female samples during the MPW (**Methods**). We revealed male-biased expression of 19 protein-coding genes, 13 of which are known androgen targets (e.g. *MAFB, CSRP2, CYP1B1, AFF3, TIPARP*)^42^ (**Figure 2e**). Seeking to determine if these genes are conserved across species, we re-analysed a murine scRNA-seq dataset of male and female external genitalia during the MPW^36^ and found that *Mafb* and *Csrp2* also show male-biased expression in the murine equivalent of the *corpus spongiosum* (**Methods, Supplementary Figure 5c-h**).

Additionally, we identified genes with sexually dimorphic expression in the human urethral epithelium. *SCGB1A* and *PTPRD,* the most upregulated genes in males (**Figure 2f**), are known for their roles in the formation of canalised epithelial structures in other tissues: *SCGB1A* has been implicated in the tubular organisation of human-derived *in vitro* bronchioids^43^, while PTPRD is recruited to epithelial adherens junctions at the time of cell-cell contact^44^.

Finally, knowing that mesenchymal differentiation directs epithelial differentiation^1^, we inferred sexually dimorphic cell-cell communication events between the human *corpus spongiosum* and the urethral epithelium using *CellphoneDB*^45^. Our analysis revealed a candidate interaction in males between *JAG1* (upregulated in the *corpus spongiosum*) and *NOTCH2/3* (expressed in the urethral epithelium), suggesting heightened NOTCH signalling (**Figure 2g**). This was further supported by the expression of downstream *HES1*. Moreover, we identified potential male-biased interactions between receptor proteins involved in adherens junctions (*NRP1, NRXN3, PTPRD*) expressed by the urethral epithelium and their ligands (*SEMA3A/PGF, NLGN1/NLGN2/NLGN3/CLSTN1*) which are upregulated in the *corpus spongiosum* (**Figure 2g**). These findings, alongside the male-specific upregulation of *SCGB1A* and *PTPRD* in the urethral epithelium, support the key role of adherens junction signalling in enabling urethral canalisation in males.

Altogether, our findings shed light on establishment of sexual dimorphism in the external genitalia by elucidating the genes and mesenchymal-epithelial interactions that likely mediate urethral canalisation in the penis (Figure 2h).

### Ontology, migration and regression of the Müllerian ducts

Unlike the external genitalia, the Müllerian and Wolffian ducts undergo significant changes in cell type composition between the sexes from 6 to 21 PCW, with sexually matched ducts differentiating into the internal genitalia and unmatched ducts regressing. The Müllerian ducts initially consist of simple meso-epithelial tubes that are specified from the extra-gonadal coelomic epithelium^46^ around 6 PCW. They migrate caudally in response to signalling from the Wolffian ducts^34^ (which emerge earlier, around 4 PCW, a developmental stage difficult to access and therefore not included in our study), and eventually fuse at the urogenital sinus (Figure 3a).

**Figure 3.**
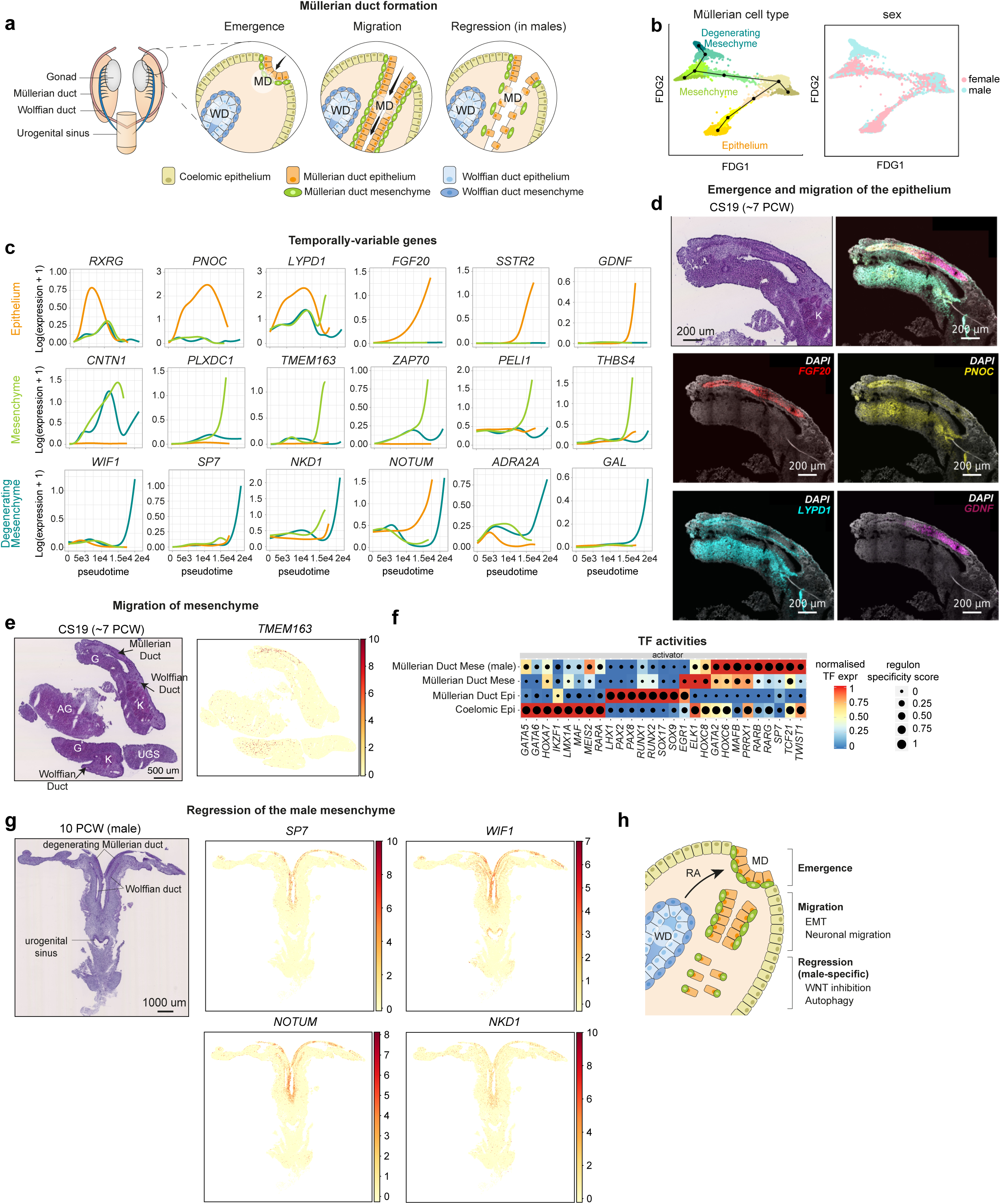
Ontology, migration and regression of the Müllerian ducts. **a,** Schematic illustration of the major steps of Müllerian duct formation (inspired by^46^) along with the cell types involved. **b,** Batch corrected force directed graph (FDG) visualisation of 6-8 PCW scRNA-seq cells from the coelomic epithelium, Müllerian duct epithelium and Müllerian duct mesenchyme coloured by cell type (left) and chromosomal sex (right). Trajectories reconstructed with *Slingshot* are overlaid on the embedding. **c,** Smoothed splines of key genes involved in each differentiation trajectory of the Müllerian duct epithelium and mesenchyme (including male degenerating mesenchyme) from the coelomic epithelium. **d,** Haematoxylin and eosin (H&E) image and high-resolution, large-area imaging of a representative section of a Carnegie Stage (CS) 19 sample with intensity proportional to smFISH signal for *FGF20* (red, migrating Müllerian duct epithelium), *PNOC* (yellow, rostral migrating Müllerian duct epithelium), *LYPD1* (cyan, rostral Müllerian duct epithelium) *GDNF* (magenta, caudal migrating Müllerian duct epithelium). **e,** H&E image and *ISS* data of a representative section of a CS19 sample showing the measured expression of *TMEM163* in the Müllerian duct mesenchyme. **f,** Heatmap showing the inferred regulon activity (from both scRNA-seq and scRNA-seq data 6-8 PCW samples) of the transcription factors (TFs) (x-axis) involved in the formation of the Müllerian duct in each relevant cell type (y-axis). The colour scale is proportional to the expression of the TF while the size of the dot reflects the importance of the TF regulon in each cell type. **g,** H&E image and *ISS* data of a representative section of a 10 PCW male sample showing the measured expression of *SP7*, *WIF1*, *NOTUM*, *NKD1* in the degenerating Müllerian duct mesenchyme. **h,** Schematic illustration summarising the putative mechanisms underpinning the formation of the human Müllerian duct as found in our analysis. AG: adrenal gland, G: gonad, K: kidney, MD: Müllerian duct, UGS: urogenital sinus; WD: Wolffian duct.

To shed light on the early events of human Müllerian duct formation, we subsetted the scRNA-seq data to the cell types that play key roles in its emergence, migration, and initial regression (between 6 and 8 PCW): the anterior mesonephric coelomic epithelium and the undifferentiated Müllerian epithelium and mesenchyme. Using *Slingshot*^47^ we recovered two trajectories from the progenitor coelomic epithelium population to the Müllerian epithelium and mesenchyme, respectively. Notably, we also found a male-specific degenerating mesenchymal lineage branching off the Müllerian mesenchyme (**Figure 3b**).

Within the Müllerian epithelial lineage, genes such as *RXRG, PNOC,* and *LYPD1* are transiently upregulated at the onset of coelomic epithelial cell differentiation (**Figure 3c**). *ALDH1A1* expressed by the Wolffian epithelium is the likely source of retinoic acid signalling via the *RXRG-RARG* axis (**Supplementary Figure 6a**). As the trajectory progresses, migratory genes such as *FGF20*, *SSTR2*, *LGI1, CALCA*, and *GDNF*, known for their roles in neuronal migration and axonal outgrowth^48–52^, become upregulated (**Figure 3c**). We validated the expression of *PNOC, LYPD1, FGF20, GDNF,* and *CALCA* by single molecule fluorescence in situ hybridization (smFISH) **(Figure 3d**, **Supplementary Figure 6b)**.

The Müllerian mesenchymal lineage, in turn, is initially characterised by upregulation of epithelial-to-mesenchymal transition (EMT) markers such as *CNTN1*^53^ (**Figure 3c**). It then upregulates migratory genes *PLXDC1*, *ZAP70*, and *TMEM163*, the latter of which we confirmed with *ISS* (**Figure 3c, e**). By contrast, the male-specific degenerating branch shows increased expression of the autophagy modulators *ADRA2A*^54^, *GAL*^55,56^ and *LAMP5*^57^, the WNT signalling inhibitors *NOTUM* and *NKD1*, as well as two previously reported markers from the mouse literature, *SP7*^58^ (whose activity was confirmed by scATAC-seq) and *WIF1*^59^ (also a WNT inhibitor) (**Figure 3c, f, Supplementary Figure 6c-f**). The cell type specificity of *NOTUM*, *NKD1*, *SP7* and *WIF1* was validated by *ISS* (**Figure 3g**).

Overall, our multi-modal approach elucidates that human Müllerian duct formation requires coordinated expression of migration genes in both mesenchyme and epithelium, alongside male-specific upregulation of WNT inhibitors and autophagy markers in the mesenchyme during Müllerian degeneration (**Figure 3h**).

### Regulators of Müllerian and Wolffian duct mesenchymal patterning

We then proceeded to investigate the differentiation and patterning of the Müllerian and Wolffian ducts into their final organ derivatives. In females, once the migration process is complete, the Müllerian ducts are regionalised along the rostro-caudal axis - cells of the distinct segments acquire different identities to form the fallopian tubes, uterus, cervix and the upper part of the vagina. Similarly, in males, the Wolffian ducts regionalise along the rostro-caudal axis, giving rise to the epididymis, vas deferens and seminal vesicle.

To explore how regional gene expression in the Müllerian and Wolffian-derived cells controls organ formation along the developing reproductive tracts, we used our spatially-resolved transcriptomics data to generate a computational representation of the rostro-caudal axis^60^ in females (from the fallopian fimbriae to the end of the Müllerian vagina^60^) and males (from the efferent ductules to the initial segment of the vas deferens) (**Figure 4a-e**, **Supplementary Figure 7a-b**, **Supplementary Note 2, 3)**. For females, where we have several *ISS* samples available, we projected the spatial axis values from *ISS* onto the scRNA-seq data, assigning pseudospace coordinates to dissociated cells (**Figure 4f**, **Supplementary Figure 7c**, **Supplementary Note 2**). Finally, we modelled gene expression changes along each axis in *ISS*-imputed scRNA-seq data (only for Müllerian ducts) and *10x Visium* (for both Müllerian and Wolffian ducts; **Supplementary Note 2, 3**).

**Figure 4.**
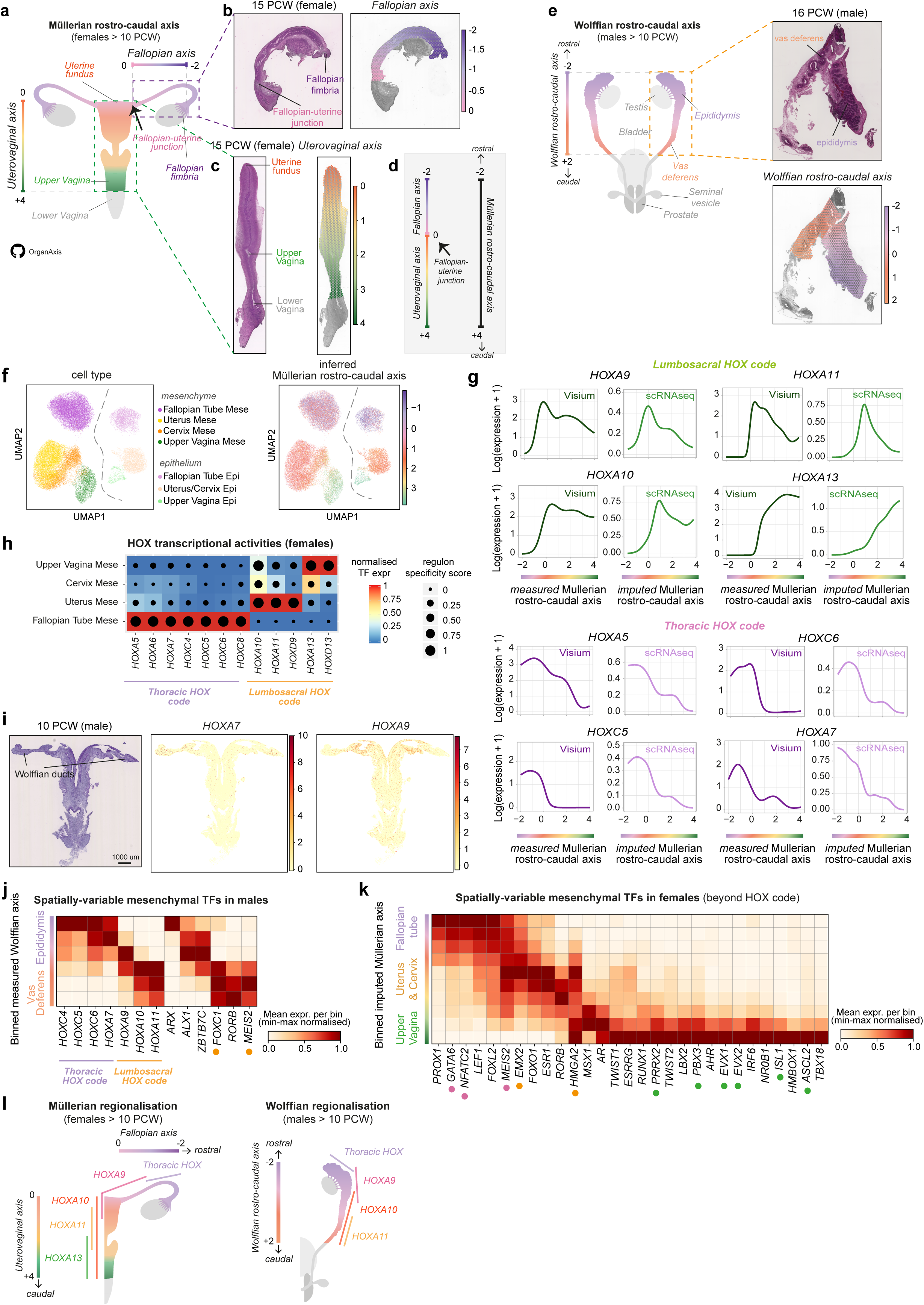
Regulators of Müllerian and Wolffian duct mesenchymal patterning. **a,** Schematic illustration of the fallopian axis (which spans the organ from the fimbriae to the fallopian-uterine junction) and the uterovaginal axis (which goes from uterine fundus until the end of the mesodermal vagina). Both axes are computed from the hematoxylin and eosin (H&E) images of *10x Visium* and *ISS* female samples >10 PCW. **b,** (left) H&E image of a representative 15 PCW fallopian tube sample profiled with *10x Visium* with annotated fimbriae and the fallopian-uterine junction. (right) Fallopian axis values in each *10x Visium* spot shown over the H&E image. **c,** (left) Stitched H&E images of two consecutive sections of a representative 15 PCW uterovaginal canal sample profiled with *10x Visium* with annotated uterine fundus, beginning of upper (Müllerian) vagina, and beginning of lower (endodermal) vagina. (right) Uterovaginal axis values in each *10x Visium* spot shown over the H&E images of the two consecutive sections. **d,** Schematic diagram of the Müllerian rostro-caudal axis derived from the concatenation of the fallopian axis and uterovaginal axis illustrated in **a**. **e,** (left) Schematic illustration of the Wolffian rostro-caudal axis, which spans the developing epididymis and vas deferens and is computed from the H&E images of *10x Visium* male samples >10 PCW. (top-right) H&E image of a representative 16 PCW male sample profiled with *10x Visium* with annotated epididymis and vas deferens. (bottom-right) Wolffian rostro-caudal axis values in each *10x Visium* spot shown over the H&E image. **f,** (left) Batch-corrected Uniform Manifold Approximation and Projection (UMAP) embedding of epithelial and mesenchymal cells (n = 57,667) in female scRNA-seq samples >10 PCW coloured by cell type. (right) Batch-corrected UMAP embedding of epithelial and mesenchymal cells in female scRNA-seq samples >10 PCW coloured by imputed Müllerian rostro-caudal axis value. **g,** Smoothed splines of *HOX* transcription factors belonging to the lumbosacral and thoracic code along the measured/imputed Müllerian rostro-caudal axis in mesenchymal spots/cells from scRNA-seq/*10x Visium* data. **h,** Inferred regulon activity (from both scRNA-seq and scATAC-seq data of female >10 PCW samples) of the *HOX* transcription factors (TFs) (x-axis) involved in patterning the differentiating Müllerian duct mesenchyme (y-axis). The colour scale is proportional to the expression of the TF while the size of the dot reflects the importance of the TF regulon in each cell type. **i,** (left) H&E image of a representative 10 PCW male sample profiled with *In Situ* Sequencing (*ISS*) alongside (right) the measured expression of *HOXA7* and *HOXA9* in the Wolffian mesenchyme. **j,** Heatmap showing the min-max normalised expression measured in *10x Visium* data of prioritised spatially variable mesenchymal TFs (x-axis) along the binned measured Wolffian rostro-caudal axis (y-axis). **k,** Heatmap showing the min-max normalised expression measured in scRNA-seq data of prioritised spatially variable mesenchymal TFs beyond the *HOX* code (x-axis) along the binned imputed Müllerian rostro-caudal axis (y-axis). **l,** Schematic illustration summarising the mesenchymal patterning by HOX code genes along the differentiating Müllerian and Wolffian ducts found by our analysis.

We first examined the expression of the four lumbosacral *HOX* genes (*HOXA9/10/11/13*) known to orchestrate the rostro-caudal regionalisation of the Müllerian and Wolffian mesenchyme in rodents^11^. In females, *HOXA10-11* was upregulated in the uterocervical and *HOXA13* in the cervicovaginal mesenchyme, respectively, aligning with previous studies^11^ (**Figure 4g**). However, while literature suggests that *Hoxa9* is upregulated throughout the fallopian tubes in mice^11^, human *HOXA9* exhibits increased expression in the caudal fallopian tube and uterocervical mesenchyme but is absent in the rostral fallopian tube mesenchyme (**Figure 4g**). This discrepancy prompted us to investigate other *HOX* genes potentially involved in patterning the rostral region of the human fallopian tubes.

Thoracic *HOX* code members (including *HOXA5*, *HOXC5*, *HOXC6*, and *HOXA7*) showed increased expression in the rostral fallopian tube mesenchyme, with a gradual decrease moving caudally (**Figure 4g**). Integrative analysis of scRNA-seq/scATAC-seq further confirmed the activity of thoracic *HOX* regulons in the fallopian tube mesenchyme, while the lumbosacral *HOX* regulons were active in the uterovaginal mesenchyme (**Figure 4h**). Moreover, the Wolffian mesenchyme, where the upper Müllerian ducts (corresponding to the region that gives rise to fallopian tubes) are embedded, appears to be patterned by *HOXA7* rostrally and *HOXA9* caudally from the earliest stages of development (**Figure 4i**). This suggests that the Müllerian duct mesenchyme adopts the *HOX* code of the adjacent Wolffian mesenchyme as it elongates beside it (**Supplementary Figure 7d**). Consistent with the Wolffian mesenchyme already being patterned early in development, in males the thoracic code later (∼10-21 PCW) marks the upper half of the epididymis while *HOXA9* is restricted to the lower half (**Figure 4j, Supplementary Figure 7e**).

We then explored the transcription factors (TFs) that likely determine the regional specificity of the mesenchyme within the ducts beyond the *HOX* code. In the Müllerian mesenchyme we revealed a decreasing gradient of *GATA6*, *PROX1*, *NFATC2*, and *FOXL2*, and an increasing gradient of *PBX3*, *PRRX2*, *EVX1*, *EVX2*, *LBX2*, *AHR*, *AR*, and *ISL1*, along the rostro-caudal axis, some of which were also found to be active by means of scATAC-seq (**Figure 4k**, **Supplementary Figure 7f,h**). These TFs include homeobox genes (*GATA6*, *PBX3*, *PRRX2*, *EVX1*, *EVX2, LBX2,* and *ISL1*) implicated in mesodermal rostro-caudal patterning of other organs and species^61–66^. Additionally, the central portion of the Müllerian axis (corresponding to the uterus and cervix) is characterised by upregulation of *EMX2*, *ESR1*, *FOXO1, MEIS2* and *RORB* (**Figure 4k**).

We observed a distinct pattern of TFs within the Wolffian-derived mesenchyme, with *ARX* and *ALX1* marking the rostral and caudal portions of the epididymis (the male analogue of the fallopian tubes), respectively (**Figure 4j**). The portion of the Wolffian axis corresponding to the upper vas deferens (the male analogue of the uterus) showed specific expression of *FOXC1*, as well as shared upregulation of *MEIS2* and *RORB* with its female uterine counterpart (**Figure 4j**, **Supplementary Figure 7g,i**). Due to damage to the vas deferens during dissections, we could not define shared and specific TFs in the lower part of the vas deferens and seminal vesicles.

Altogether, our work has led to refinement of the *HOX* code responsible for mesenchymal regionalisation in the differentiating human Müllerian and Wolffian ducts (**Figure 4l**). Moreover, we have identified novel spatially-variable TFs, with some being shared between the female and male ducts and others unique to each.

### Cell-cell communication along the Müllerian and Wolffian duct niches

Heterotypic co-culturing of epithelial and mesenchymal cells of the reproductive tract showed that mesenchymal cells in the ducts first acquire their regional identity and then instruct the adjacent epithelium to differentiate accordingly^1^. We next performed cell-cell communication analysis to identify the specific interactions between the mesenchyme and epithelium along the Müllerian and Wolffian duct axis (Figure 5a, Supplementary Figure 8a-l).

**Figure 5.**
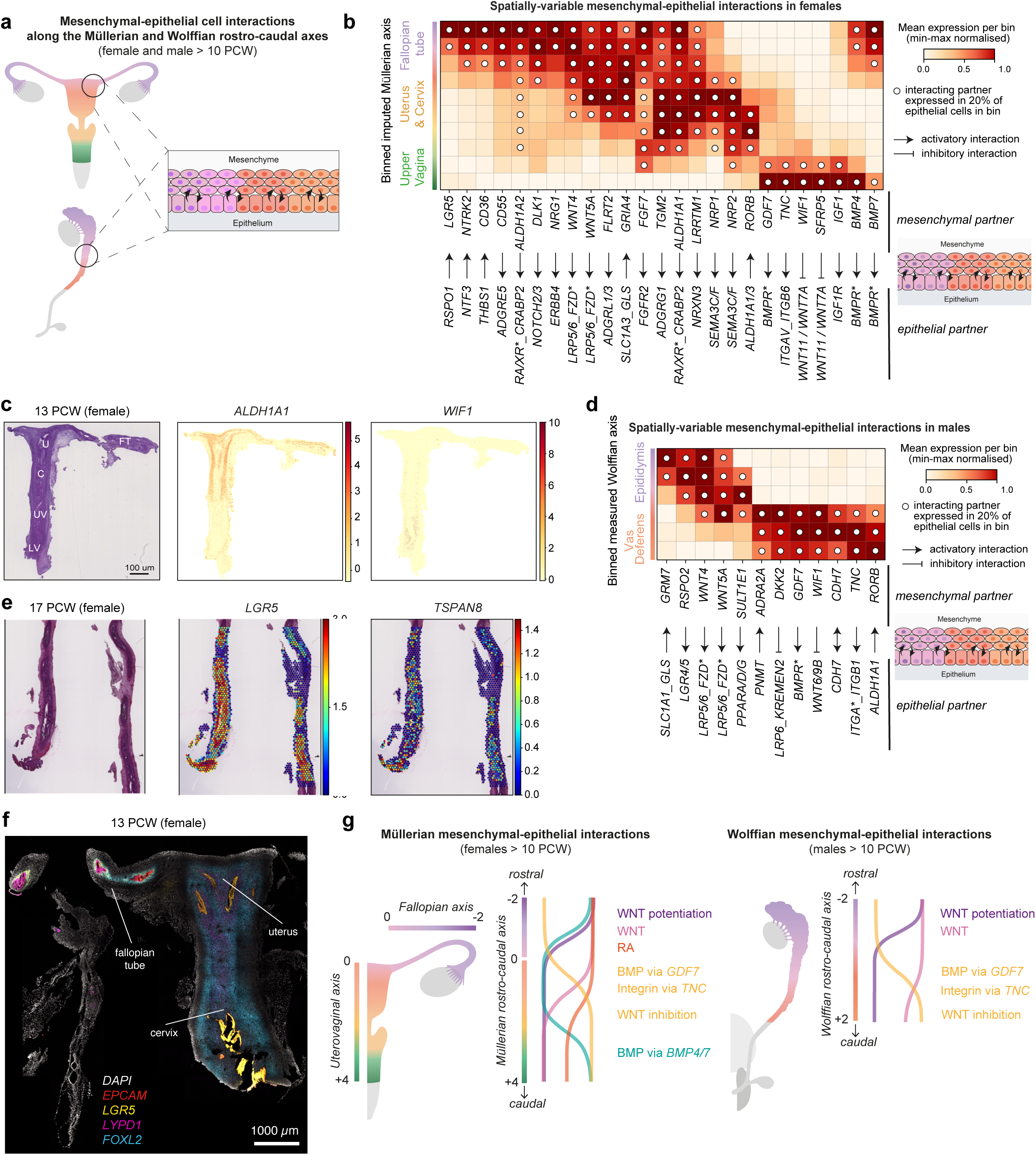
Cell-cell communication along the Müllerian and Wolffian duct niches. **a,** Schematic illustration showing the bidirectional cell-cell communication between the mesenchyme and epithelium of the differentiating Müllerian and Wolffian ducts along their rostro-caudal axes in female and male samples >10 PCW. **b,** Heatmap showing the min-max normalised expression measured in scRNA-seq data of prioritised spatially variable mesenchymal ligands or receptors (x-axis) along the binned imputed Müllerian rostro-caudal axis (y-axis). Interacting partners for each spatially variable mesenchymal ligand/receptor are reported on the x-axis below the activatory or inhibitory arrows. A dot in a bin indicates that the interacting partner of the spatially variable mesenchymal ligand/receptor is expressed in at least 20% of epithelial cells in that bin. **c,** (left) Hematoxylin and eosin (H&E) image of a representative 13 PCW female sample profiled with *In Situ* Sequencing (*ISS*) alongside (right) measured expression of *ALDH1A1* and *WIF1.* **d,** (left) H&E image of a representative 17 PCW fallopian tube sample profiled with *10x Visium* alongside (right) measured expression of *LGR5* and *TSPAN8* in each spot. **e,** Heatmap showing the min-max normalised expression measured in *10x Visium* data of prioritised spatially variable mesenchymal ligands or receptors (x-axis) along the binned measured Wolffian rostro-caudal axis (y-axis). Interacting partners for each spatially variable mesenchymal ligand/receptor are reported on the x-axis below the activatory or inhibitory arrows. A dot in a bin indicates that the interacting partner of the spatially variable mesenchymal ligand/receptor is expressed in at least 20% of epithelial cells in that bin. **f,** High-resolution, large-area imaging of a representative section of a 13 PCW female sample with intensity proportional to smFISH signal for *EPCAM* (red, epithelium), *LGR5* (yellow, fallopian tube mesenchyme and uterocervical epithelium), *LYPD1* (magenta, rostral fallopian tube epithelium), *FOXL2* (cyan, fallopian and uterocervical mesenchyme). **g,** Schematic illustration summarising the mesenchymal-epithelial signalling present along the differentiating Müllerian and Wolffian ducts found in our analysis. C: cervix; FT: fallopian tube; LV: lower (endodermal) vagina; UT: uterus; UV: upper (Müllerian) vagina.

We found heightened activity of *WNT* and retinoic acid signalling (mediated by mesenchymal-expressed ligands *WNT4/WNT5A* and *ALDH1A1*) in the fallopian tubes and uterus (**Figure 5b-c**). This is opposed by an increasing gradient of *WNT* inhibition (driven by *WIF1* and *SFRP5* upregulated in the mesenchyme) in the upper vagina, corroborating existing mouse literature^67,68^ (**Figure 5b-c**). Similarly, along the Wolffian duct axis, we observed opposing patterns of WNT activation and inhibition between the rostral and caudal segments, coupled with potentiation of WNT signalling mediated by *RSPO2*-*LGR5* in the upper epididymis (**Figure 5d**).

In the upper vagina, we observed increased signalling through the *IGF1-IGF1R* axis, integrin pathways involving *TNC*, and *BMP* activity mediated by *GDF7* and *BMP4/7* through *BMPR*, with each respective ligand being expressed in the mesenchyme*. BMP* signalling could induce the upregulation of *RUNX1* and *TP63* in the adjacent epithelium, as previously reported in mice^69^ and in keeping with our analysis of spatially variable TFs in the differentiating Müllerian epithelium (**Figure 5b**, **Supplementary Figure 8d, i-j**). While *GDF7* and *TNC* expression peaked in the upper vagina, they were already expressed in the uterocervical mesenchyme (**Figure 5b**). This expression pattern was mirrored in males, where *BMP* signalling via *GDF7* and integrin signalling via *TNC* (both mesenchymal-expressed) were upregulated in the initial segment of the vas deferens (**Figure 5d)**.

Signalling between mesenchyme and epithelium during ductal regionalisation can be bi-directional, with signals from the epithelium also influencing the fate of the mesenchyme^1^. Consistent with this, the fallopian tube mesenchyme upregulated the *LGR5* receptor, which could potentially receive signals from the adjacent epithelium expressing its cognate ligand *RSPO1* (**Figure 5b**). The interaction of *RSPO1* with *LGR5*, known to enhance canonical *WNT* signalling in other systems, along with the co-expression of *TSPAN8* in the fallopian mesenchyme, suggests characteristics reminiscent of a stem cell niche (**Figure 5e-f**, **Supplementary Figure 8b**).

In summary, by investigating mesenchymal-epithelial cell interactions along the Müllerian and Wolffian rostro-caudal axes, we identified shared and sex-specific cell communication events that are likely pivotal in determining epithelial identity during the regionalisation of the reproductive ducts (**Figure 5g**).

### Regionalisation of the fallopian tube and epididymis occurs during fetal development

In adulthood, the non-ciliated epithelia of the fallopian tubes and epididymis are functionally regionalised to support sperm maturation – reflected in marked differences in gene expression – yet it is unclear if and when this regional differentiation occurs during fetal development^70^. To evaluate the *in utero* transcriptional gradients at the genome-wide level, we leveraged our Müllerian and Wolffian rostro-caudal axes analysis framework from the previous section. Here, we focused on the rostral portion of each axis, examining intra-organ gene expression changes within the developing fallopian tube and epididymis epithelia in samples between 10 and 21 PCW (**Figure 6a-b**, **Supplementary Note 4**).

**Figure 6.**
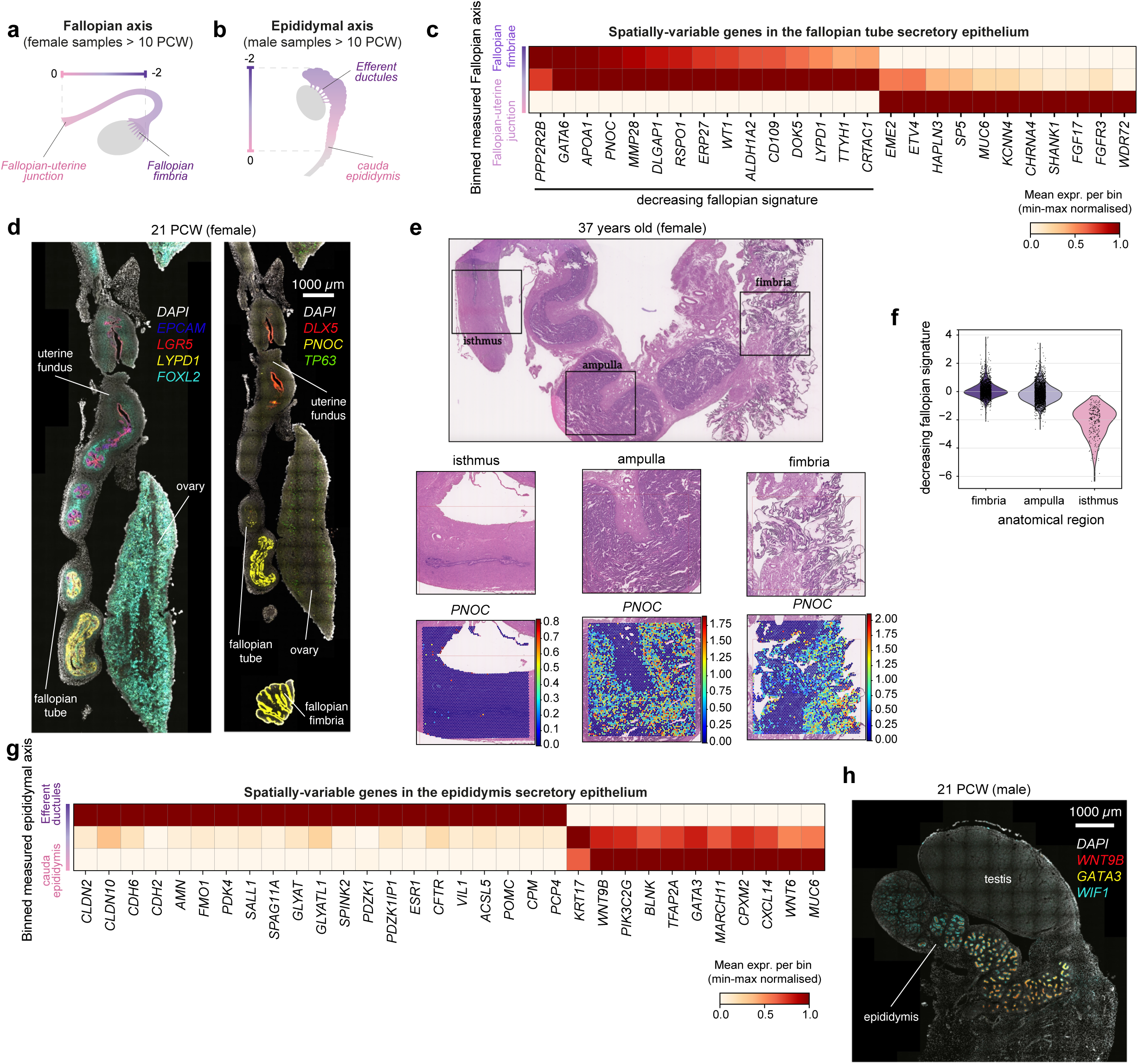
Regionalisation of the fallopian tube and epididymis occurs during fetal development. **a,** Schematic illustration of the derivation of the fallopian axis in female samples >10 PCW (this is part of the Müllerian rostro-caudal axis shown in Figure 4). **b,** Schematic illustration of the derivation of the epididymal axis in male samples >10 PCW (this is part of the Wolffian rostro-caudal axis shown in Figure 4). **c,** Heatmap showing the min-max normalised expression measured in *10x Visium* data of spatially variable non-ciliated epithelial genes (x-axis) along the binned fallopian axis (y-axis). **d,** (left) High-resolution, large-area imaging of a representative section of a 21 PCW female sample with intensity proportional to smFISH signal for *EPCAM* (blue, epithelium), *LGR5* (red, fallopian tube mesenchyme and uterine epithelium), *LYPD1* (yellow, rostral fallopian tube epithelium), *FOXL2* (cyan, fallopian tube and uterine mesenchyme). (right) High-resolution, large-area imaging of a representative section of a 21 PCW female sample with intensity proportional to smFISH signal for *DLX5* (red, uterine epithelium), *PNOC* (yellow, rostral fallopian tube epithelium), *TP63* (green, oocytes). **e,** (top) H&E image of a 37 years old fallopian tube sample with highlighted regions that were placed onto three different capture areas for spatial transcriptomics profiling by *10x Visium*. (middle) H&E images of the three capture areas highlighted in (top). (bottom) Normalised, log-transformed expression of *PNOC* in each *10x Visium* spot of the three capture areas. **f,** Violin plot showing the decreasing signature score (of the genes plotted in panel **c**) in the epithelial *10x Visium* spots of each of the three anatomical regions. The significance of the decreasing trend was evaluated with Jonckheere’s trend test (p-value = 5e04). **g,** Heatmap showing the min-max normalised expression measured in *10x Visium* data of spatially variable non-ciliated epithelial genes (x-axis) along the binned epididymal axis (y-axis). **h,** High-resolution, large-area imaging of a representative section of a 21 PCW male sample with intensity proportional to smFISH signal for *WNT9B* (red, caudal epididymal epithelium), *GATA3* (yellow, caudal epididymal epithelium), *WIF1* (cyan, epididymal mesenchyme).

In the non-ciliated epithelial cells in the fallopian tube, we identified a set of 15 genes (including *PNOC*, *LYPD1*, *APOA1*, *WT1*, *GATA6, ERP27*, and *CRTAC1*) whose expression decreases rostro-caudally from the developing fimbria to the isthmus **Figure 6c**, **Supplementary Figure 9a**). Notably, *PNOC* and *LYPD1* expression is already restricted to the rostral portion of the epithelium during Müllerian duct emergence, indicating that some degree of regionalisation begins early in development (**Figure 3d**). Additionally, we observed upregulation of *MUC6*, *WDR72* and *KCNN4* in the fallopian isthmus (**Figure 6c**). Orthologues of these genes are involved in isthmus-specific epithelial secretions in other species^71,72^.

It is known that some genes change their expression^73,74^ along the rostrocaudal axis of the fallopian tubes in adults, yet a comprehensive study is missing. Thus, to determine if the spatial gradient we identified in the fetus persists into adulthood, we generated *10x Visium* spatial transcriptomics data from three regions of an adult fallopian tube (fimbria, ampulla, and isthmus) and scored the epithelial spots in each regions for the average expression of this gene set (**Figure 6e**). The decreasing rostro-caudal trend of the genes identified during fetal development is indeed preserved in adulthood (**Figure 6f**). Additionally, as in the fetal fallopian tubes, we observed upregulation of *MUC6*, *WDR72* and *KCNN4* in the adult fallopian isthmus (**Supplementary Figure 9b-c**).

Within the non-ciliated epithelium of the fetal epididymis, we uncovered genes with a rostral-bias, including *ESR1*, *SALL1*, *VIL1*, *CFTR, SPAG11A, PDZK1,* and *GLYAT*, which are known adult regulators implicated in fluid reabsorption and sperm maturation^75–78^ (**Figure 6g**, **Supplementary Figure 9d**, **Supplementary Figure 8k-l**). Additionally, cell adhesion genes like claudins (*CLDN2/10*) and cadherins (*CDH2/6*) were enriched in the rostral portion of the epididymis, consistent with findings in adults^76^. Conversely, several genes whose expression increased rostro-caudally – *GATA3, WNT9B, TFAP2A*, *CXCL14, MARCH11, CPXM2,* and *BLNK* – have also been reported in adults and are associated with immune response regulation^76^ (**Figure 6g-h**, **Supplementary Figure 8k-l**).

Taken together, our findings revealed that the regional differentiation of the human fallopian tube and epididymis begins during fetal development.

### Mapping potential disruptions to reproductive tract development

Exogenous agents, including drugs and environmental chemicals, can disrupt developmental programs *in utero*, and manifest as reproductive disorders later in life^79,80^. We investigated what reproductive cell types can potentially be targeted by known drugs that can cross the maternal-fetal interface using *drug2cell*^81^, a computational drug screening tool that utilises known drug-protein interactions to predict cellular targets. We focused particularly on drugs with potential to disrupt reproductive epithelia, as this cellular compartment is frequently implicated in disease, and identified 47 such compounds (with known ATC classification^82^) (**Figure 7a**, **Supplementary Table 10**).

**Figure 7.**
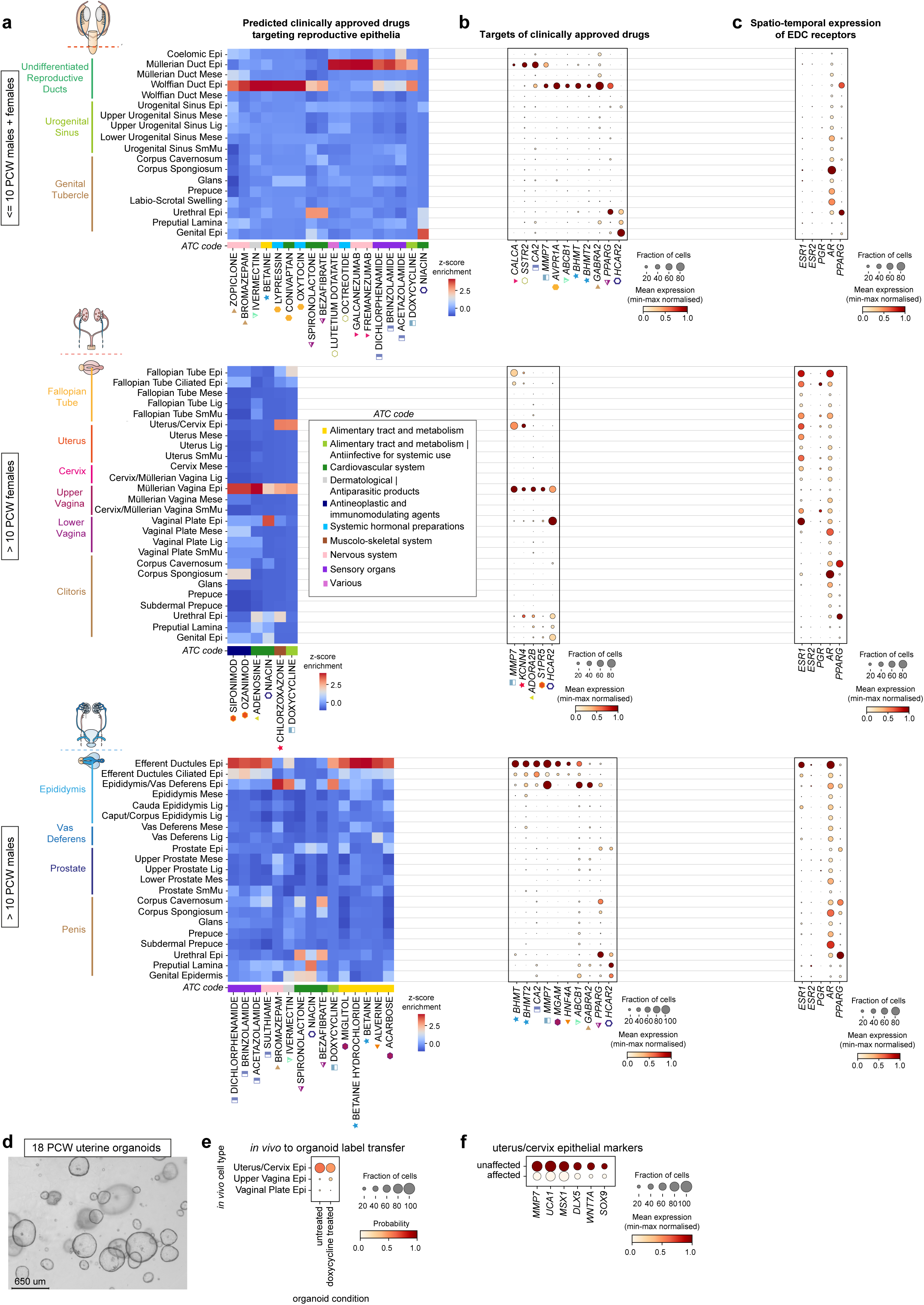
Mapping potential disruptions to reproductive tract development. **a,** Heatmap showing the z-score enrichment of targets of clinically approved drugs (x-axis) found to be specifically impacting the epithelial compartment of reproductive tract organs among reproductive-specific cell types (y-axis) identified in our scRNA-seq dataset. **b,** Dot plot showing the log-transformed, min-max normalised expression of the target genes (x-axis) of the clinically approved drugs in **a,** in each reproductive-specific cell type (y-axis) identified in our scRNA-seq dataset. Corresponding drugs and targets are indicated with the same symbol. **c,** Dot plot showing the log-transformed, variance-scaled expression of the steroidogenic receptors (x-axis) whose activity can be disrupted by endocrine disrupting chemicals (EDCs) in each reproductive-specific cell type (y-axis) identified in our scRNA-seq dataset. For panels **a**, **b**, and **c**, the visualisations are split in <=10 PCW female and male samples (top); >10 PCW female samples (middle); >10 PCW male samples (bottom). **d,** Bright-field microscopy image of uterine epithelial organoids derived from an 18 PCW female sample at day 2 of doxycycline treatment. **e,** Dot plot showing the predicted probability from each epithelial *in vivo* cell type in the 18 PCW female sample from which the organoids were derived and the two organoid libraries (untreated and treated with doxycycline). **f,** Dot plot showing the log-transformed, variance-scaled expression of the uterine epithelial markers in the “unaffected” and “affected” cells of the organoid library treated with doxycycline.

The targets of monoclonal antibodies like fremanezumab (targeting *CALCA*) and somatostatin agonists such as octreotide (binding to *SSTR2*), which are used in the treatment of migraines and neuroendocrine tumours respectively, are upregulated in the migratory Müllerian epithelium relative to all other cell types (**Figure 7a-b**). Conversely, the early Wolffian duct epithelium could be specifically targeted by drugs like conivaptan (acting on *AVPR1A*) and spironolactone (targeting *PPARG*) which are used for type 1 and 2 diabetes respectively (**Figure 7a-b**). Both Müllerian and Wolffian duct epithelia, as well as their derivatives, particularly the uterocervical and vas deferens epithelia, also exhibit susceptibility to the antibiotic doxycycline (binding to *MMP7*) (**Figure 7a-b**). As a proof-of-principle experiment, we tested the predicted disruption by doxycycline using a uterine epithelial organoid line treated with doxycycline, and observed a reduction in the expression of uterine epithelial markers (*UCA1, MSX1*, *DLX5, WNT7A, SOX9)*, as well as *MMP7* as expected, in approximately 33% of treated cells, indicating an impact on cell identity (**Figure 7d-f**).

We also assessed the potential impact of endocrine-disrupting chemicals commonly found in substances such as plastics, and evaluated the dynamics of steroidogenic hormone receptor expression throughout gestation. Receptors for estrogen (*ESR1/2*) and progesterone (*PGR*), whose activities are affected by bisphenol A^83^; androgen receptor (*AR*), disrupted by phthalate esters^84,85^; and peroxisome proliferator-activated receptor gamma (*PPARG*), recently identified as a target of per- and polyfluoroalkyl substances (PFAS)^86^, were all expressed in both male and female reproductive tracts. Before 10 PCW, *ESR1/2* and *PGR* were not detected (**Figure 7c**).

After 10 PCW, *ESR1* was upregulated in most Müllerian duct derivatives, in the vaginal plate epithelium, and in the caput epididymis epithelium, while *PGR* was lowly expressed in the ciliated fallopian tube epithelium and in the smooth muscle of the upper vagina (**Figure 7c**). *AR* was highly expressed in the fallopian tube epithelium, the vaginal plate mesenchyme, and the epithelium of the entire epididymis after 10 PCW (**Figure 7c**). Upregulation of *AR* was also observed in the urogenital sinus and genital tubercle derivatives throughout gestation. *PPARG* was upregulated in the epithelial cells of the Wolffian duct early in development and later in the caudal epididymis and vas deferens, as well as in the urethral epithelium across all stages (**Figure 7c**).

In summary, our integrated atlas allowed us to predict when and where external agents, such as clinically approved drugs and endocrine-disruptive chemicals, have the potential to act during gestation.

## Discussion

Congenital reproductive tract disorders affect more than 3% of female^19^ and 0.8% of male^20^ newborns, and yet our understanding of prenatal reproductive tract development remains limited. In this study, we generated a roadmap of the male and female reproductive tracts during key periods of sexual differentiation, detailing the temporal and spatial distribution of 52 reproductive tract-specific cell types in 80 human samples during prenatal development (6-21 PCW). We uncovered sex-specific cues driving the divergent development of reproductive organs and the selective regression of sexually unmatched ducts. In addition, by capturing the evolving complexity of epithelial and surrounding mesenchymal compartments from progenitor to differentiated states, we provided cellular and molecular insights into the organisation of the developing tract, whereby early axial gradients are translated into defined cell lineages and distinct tissue structures.

To investigate sexual dimorphism in the external genitalia we focused on the downstream genes regulating androgen-driven urethral canalisation in males, a process disrupted in hypospadias^9,25^. We found male-biased gene expression in both the mesenchymal *corpus spongiosum* and urethral epithelium, as well as sexually dimorphic interactions between these cell types. Among the male-biased genes in the *corpus spongiosum*, we identified *MAFB*, which is the only mesenchymal, androgen-responsive gene essential for urethral canalisation that has been validated with a functional study in mice^40^. While surgical reconstruction of the urethra remains the primary treatment for hypospadias, its success in severe cases is often limited by insufficient graft material. Our insights into the molecular drivers of urethral canalisation could guide tissue engineering efforts to develop more effective graft materials^87,88^. Additionally, we identified novel male-specific genes responsible for Müllerian duct regression, reinforcing the role of WNT inhibitors as reported in murine functional studies^17,59^ and providing a molecular foundation for autophagy as an underlying mechanism in this process, a hypothesis previously suggested based on electron microscopy observations^89^.

Following the regression of the unmatched ducts, the differentiation of sex-specific organs from the remaining ducts depends on the precise establishment of spatial boundaries between the organs. To determine what TFs and cell-cell communication events drive ductal regionalisation in humans, we computationally derived Müllerian and Wolffian rostro-caudal axes using our spatial transcriptomics data and adapting the OrganAxis framework^60^. This approach was needed as subtle differences in gene expression were not detectable in single-cell transcriptomic data of dissociated samples and the anatomical intra-organ regionalisation was not yet evident at these developmental stages. Our analysis revealed that thoracic *HOX* genes are expressed in the upper fallopian tubes and epididymis, challenging the traditional view that mesenchymal regionalisation is governed solely by lumbosacral *HOX* genes. Moreover, we uncovered novel TFs regulating mesenchymal differentiation, some of which are shared between Müllerian and Wolffian-derived mesenchymal cells (e.g., *MEIS2* and *RORB* in the uterus and upper vas deferens), while others are specific to one sex (e.g. *GATA6/PROX1* in the fallopian tube and *ARX1/ALX1* in the epididymis). This finding suggests that female and male organs at analogous rostro-caudal positions share common regulatory programs, while their specific roles demand unique transcriptional regulators, though further research is needed to fully understand the interplay between these mechanisms. Our comparison, however, is also limited by the inability to fully reconstruct the Wolffian duct axis due to damage to the vas deferens during dissections.

We also discovered key paracrine mesenchymal-derived signals along the Müllerian and Wolffian duct axes. For instance, *BMP4/7* are spatially confined to the upper vagina, whereas *GDF7* is more broadly expressed in the uterocervix and upper vagina, indicating distinct roles for *BMP* ligands across the female reproductive tract. This aligns with observations in the adult endometrium, where *GDF7*, but not *BMP4/7*, signalling is present^90^. Our findings can be incorporated into emerging *in vitro* models of the female and male reproductive tracts^91,92^, which are particularly needed due to differences in gene expression between murine and human reproductive organs, as well as the significant delay in advancing reproductive models compared to other bodily systems.

Furthermore, we identified the genes responsible for regionalisation in the fetal fallopian tube and epididymal non-ciliated epithelium, revealing that gene expression gradients crucial for sperm maturation in adulthood are already established prenatally. This contrasts with the uterocervical epithelium, which remains transcriptionally homogeneous until at least 21 PCW, suggesting that its adult structure may depend on hormonal influences later in life. While intra-organ regionalisation has been well-documented in tissues like the gut^93^ and liver^94,95^, and there is evidence suggesting that this process begins during fetal development in mouse gut^96^, it has not been extensively explored in female reproductive tissues, especially in the fallopian tubes. Our findings not only enhance our understanding of basic biology but also have implications for fertility conditions affecting the fallopian tubes that may be determined *in utero*. Moreover, among the transcriptomic signatures upregulated in both fetal and adult fallopian fimbriae, we uncovered genes such as *PNOC*, *APOA1*, *GATA6*, *WT1*, which are diagnostic markers and therapeutic targets of high-grade serous ovarian cancer (HGSOC)^97–99^, a malignancy known to originate in the adult fimbriae^100^. This highlights the potential of utilising fetal data, where continuous spatial modelling of gene expression is feasible due to reduced sample dimensions, to uncover new genes implicated in HGSOC.

Lastly, we used our atlas to assess *in silico* the potential toxicity of endocrine disruptors from everyday products and commonly prescribed drugs on reproductive development, where many congenital abnormalities remain unexplained. *ESR1* was upregulated in the human vaginal plate and caput epididymis epithelia, consistently with the known trans-generational effects of its agonist bisphenol A, which impair vaginal and epididymal development in rodents^83,101^. Similarly, we observed elevated AR expression throughout the epididymis and derivatives of the urogenital sinus and genital tubercle, mirroring reports of increased incidence of epididymal agenesis and hypospadias in rats exposed to phthalate esters *in utero*^102^. Among the clinically approved drugs that specifically affect reproductive epithelia, we identified diabetes medications and antibiotics like doxycycline. As a proof-of-concept, we revealed that doxycycline disrupts the expression of uterine epithelial markers in fetal organoid cultures, underscoring the utility of our atlas in predicting potential vulnerabilities of reproductive tract cell types to clinically approved drugs. Future work combining organoid models with functional assays is needed to validate these predictions and reveal the full extent of their impact on reproductive development.

In summary, our findings provide a foundational framework for understanding the signalling cues, rostro-caudal positional identity, and gene expression dynamics that guide the lineage-specification of reproductive tract tissues in both females and males. With this resource, researchers can contextualise known genetic variants linked to reproductive diseases by identifying when and in which cell types genes are expressed or chromatin regions are open. Moreover, our work paves the way for developing more complex *in vitro* models that replicate *in vivo* development, facilitating the study of disease-causing perturbations.

## Extended data figure legends

**Extended Data Figure 1. Cell type characterisation of female and male reproductive tract samples <=10 PCW. a**, Schematic representation of the computational workflow used to analyse scRNA-seq data. **b,** Batch-corrected Uniform Manifold Approximation and Projection (UMAP) embedding of female and male samples <= 10 PCW profiled with scRNA-seq (n = 177,319 cells) coloured by stage (measured in PCW), donor, sex and cell type. **c,** Dot plot showing the variance-scaled, log-transformed expression of genes (x-axis) characteristic of the annotated cell types (y-axis) detected in female and male samples <=10 PCW. Top-layer groups marker genes by developing organs. **d,** Schematic representation of the computational workflow used to analyse *In Situ* Sequencing (*ISS*) data. **e,** (top) Hematoxylin and eosin (H&E) stained image of a representative Carnagie stage (CS) 19 sample profiled with *ISS* alongside annotations of the anatomical structures using *TissueTag*^60^ and inferred cell type labels for selected cell types from the scRNA-seq dataset in the *ISS* slide using a *iss-patcher*^103^. (bottom) H&E stained image of a representative CS18 sample profiled with *ISS* alongside annotations of the anatomical structures using *TissueTag*^60^ and inferred cell type labels for selected cell types from the scRNA-seq dataset in the *ISS* slide using a *iss-patcher*^103^. AG: adrenal gland, G: gonad, K: kidney, RD: reproductive ducts, UGS: urogenital sinus.

**Extended Data Figure 2. Cell type characterisation of female reproductive tract samples >10 PCW. a,** Batch corrected Uniform Manifold Approximation and Projection (UMAP) embedding of female samples >10 PCW profiled with scRNA-seq (n = 210,810 cells) coloured by stage (measured in PCW), donor and cell type. **b,** Dot plot showing the variance-scaled, log-transformed expression of genes (x-axis) characteristic of the annotated cell types (y-axis) detected in female samples >10 PCW. Top-layer groups marker genes by developing organs. **c,** Schematic representation of the computational workflow used to analyse *10x Visium* data. **d,** Heatmap showing the z-score enrichment of each epithelial, mesenchymal, smooth muscle and ligament cell type annotated in female scRNA-seq samples >10 PCW (x-axis) in *10x Visium* spots corresponding to each histological annotation (y-axis). **e,** Hematoxylin and eosin (H&E) stained image of a representative section of a 17 PCW female sample profiled with *In Situ* Sequencing (*ISS*) alongside annotations of the anatomical structures using *TissueTag*^60^ and inferred cell type labels for selected cell types from the scRNA-seq dataset in the *ISS* slide using a *iss-patcher*^103^. **f,** *ISS* data of a representative section of a 17 PCW female sample showing the measured expression of *MYH11* (smooth muscle marker), *TP63* (vaginal stratified epithelium marker), *DLX5* (uterine epithelium marker). CE: cervix; FT: fallopian tube; UT: uterus; VA: vagina.

**Extended Data Figure 3. Cell type characterisation of male reproductive tract samples >10 PCW. a,** Batch corrected Uniform Manifold Approximation and Projection (UMAP) embedding of male samples >10 PCW profiled with scRNA-seq (n = 97,065 cells) coloured by stage (measured in PCW), donor and cell type. **b,** Dot plot showing the variance-scaled, log-transformed expression of genes (x-axis) characteristic of the annotated cell types (y-axis) detected in male samples >10 PCW. Top-layer groups marker genes by developing organs. **c,** Hematoxylin and eosin (H&E) stained image of a representative section of a 16 PCW male sample profiled with *In Situ* Sequencing (*ISS*) alongside annotations of the anatomical structures using *TissueTag*^60^ and inferred cell type labels for selected cell types from the scRNA-seq dataset in the *ISS* slide using a *iss-patcher*^103^. **d,** *ISS* data of a representative section of a 16 PCW male sample showing the measured expression of *MYH11* (smooth muscle marker), *LEFTY1* (epididymis and vas deferens epithelium marker). **e,** (top) Area plot showing the temporal distribution of cell type proportions in the internal genitalia (derived from the Müllerian ducts and urogenital sinus) from all female samples (6-21 PCW). (bottom) Area plot showing the temporal distribution of cell type proportions in the internal genitalia (derived from the Wolffian ducts and urogenital sinus) from all male samples (6-21 PCW). EP: epididymis; UGS: urogenital sinus; VD: vas deferens.

**Extended Data Figure 4. Spatial mapping of female and male external genitalia. a,** Hematoxylin and eosin (H&E) stained image of a representative section of a 14 PCW male external genitalia sample profiled with *10x Visium* along side spatial mapping of selected mesenchymal and epithelial cell types from the scRNA-seq dataset onto the corresponding *10x Visium* slide using *cell2location*^104^. Estimated cell type abundance (colour intensity) in each *10x Visium* spot is shown over the H&E image. **b,** *10x Visium* data of a representative section of a 14 PCW male external genitalia sample showing the measured expression of *FOXA1* (endodermal-derived urethral epithelium) and *KRT14* (ectodermal-derived genital skin epithelium). **c,** H&E stained image of a representative section of a 13 PCW female external genitalia sample profiled with *In Situ* Sequencing (*ISS*) alongside inferred cell type labels for selected cell types from the scRNA-seq dataset in the *ISS* slide using a *iss-patcher*^103^. **d,** *ISS* data of a representative section of a 13 PCW female external genitalia sample showing the measured expression of *FOXA1* (endodermal-derived urethral epithelium) and *KRT14* (ectodermal-derived genital skin epithelium).

**Extended Data Figure 5. Sexual dimorphism in the external genitalia and comparison with mouse. a,** Batch corrected t-distributed Stochastic Neighbour Embedding (t-SNE) embedding of downsampled 8-14 PCW scATAC-seq cells (n = 6258) from the male external genitalia coloured by cell type and developmental stage (measured in PCW). **b,** Heatmap showing the inferred regulon activity (from both scRNA-seq and scRNA-seq data 8-14 PCW male samples) of the transcription factors (x-axis) involved in the establishment of cellular diversity in the external genitalia (y-axis). The colour scale is proportional to the expression of the TF while the size of the dot reflects the importance of the TF regulon in each cell type. **c,** Schematic representation of the computational workflow used to compare human and mouse external genitalia during the masculinisation programming window. **d,** Sankey plot representing the matching neighbourhoods between the mouse and human external genitalia labelled by the most abundant cell type in the neighbourhood. **e,** Batch corrected UMAP embedding of embryonic day (E) 14.5, E16.5, E18.5 (corresponding to the masculinisation programming window in mice) scRNA-seq cells from the mouse external genitalia^36^ coloured by cell type. The cell type annotations are taken from the authors of the original study. **f,** Batch corrected UMAP embedding of embryonic day (E) 14.5, E16.5, E18.5 scRNA-seq cells from the mouse external genitalia highlighting the neighbourhoods that match the human *corpus spongiosum*. **g,** Batch corrected UMAP embedding of embryonic day (E) 14.5, E16.5, E18.5 scRNA-seq cells from the mouse external genitalia showing the expression of *corpus spongiosum* markers *FOXF1*, *SALL1*, *FOXL2*. **h,** Volcano plot showing the log fold-change (x-axis) and adjusted p-value (y-axis) of the differential expression test (*pyDESEQ2*^105^: adjusted p-value = 0.05, |logFC| > 1.25) between male and female samples within the mouse *corpus spongiosum*. Only upregulated genes in males are labelled in the plot.

**Extended Data Figure 6. Ontology, migration and regression of the Müllerian duct. a,** Dotplot showing the log-transformed, min-max normalised expression of Wolffian ligands and Müllerian receptors (y-axis) in each cell type (x-axis). Interacting ligands and receptors are connected with an arrow. **b,** Haematoxylin and eosin (H&E) stained image and high-resolution, large-area imaging of a representative section of a Carnegie Stage (CS) 23 sample with intensity proportional to smFISH signal for *KLK11* (green, coelomic epithelium), *FGF20* (red, migrating Müllerian duct epithelium), *GDNF* (cyan, caudal migrating Müllerian duct epithelium), *CALCA* (yellow, caudal migrating Müllerian duct epithelium). **c,** Schematic representation of the computational workflow used to analyse scATAC-seq data. **d,** Schematic representation of the computational workflow used to infer transcription factor activities from scRNA-seq and scATAC-seq with *SCENIC+*^106^. **e,** Batch corrected t-distributed Stochastic Neighbour Embedding (t-SNE) embedding of downsampled 6-8 PCW scATAC-seq cells (n = 1220) from the coelomic epithelium, Müllerian duct epithelium and Müllerian duct mesenchyme coloured by developmental stage (measured in PCW) and cell type. **f,** H&E image and *In Situ* Sequencing (*ISS*) data of a representative section of a 10 PCW male sample showing the measured expression of *WNT7A* and *WNT9B* in the Müllerian duct epithelium and Wolffian duct epithelium, respectively. AG: adrenal gland, G: gonad, K: kidney, UGS: urogenital sinus.

**Extended Data Figure 7. Construction of the Müllerian and Wolffian rostro-caudal axes and redefinition of the human *HOX* code. a,** Schematic representation of the computational workflow used to construct the Müllerian rostro-caudal axis from *10x Visium* and *In Situ* Sequencing (*ISS*) female samples > 10 PCW. **b,** Schematic representation of the computational workflow used to construct the Wolffian rostro-caudal axis from *10x Visium* male samples > 10 PCW. **c,** Schematic representation of the computational workflow used to impute the Müllerian rostro-caudal axis in scRNA-seq data from *ISS* female samples > 10 PCW based on a shared k-nearest neighbours embedding (approach adapted from^103^). **d,** Hematoxylin and eosin (H&E) stained image and *ISS* data of a representative section of a 10 PCW male sample showing the measured expression of *PLAC1* (expressed in the Wolffian mesenchyme) and *LHX1* (expressed in the Müllerian and Wolffian epithelia). **e,** H&E stained image and *10x Visium* data of a representative section of a 14 PCW male sample showing the measured expression of *HOXC5*, *HOXA7*, *HOXA9*, *HOXA10*, *HOXA11*. **f,** Batch corrected, t-distributed Stochastic Neighbour Embedding (t-SNE) of downsampled Müllerian-derived mesenchymal cells from female samples >10 PCW profiled with scATAC-seq coloured by stage (measured in PCW) and cell type. **g,** Batch corrected, t-SNE of downsampled Wolffian-derived mesenchymal cells from male samples >10 PCW profiled with scATAC-seq coloured by stage (measured in PCW) and cell type. **h,** Heatmap showing the inferred regulon activity (from both scRNA-seq and scRNA-seq data >10 PCW female samples) of the transcription factors (x-axis) involved in the establishment of cellular diversity in the Müllerian duct-derived mesenchymal cells (y-axis). The colour scale is proportional to the expression of the TF while the size of the dot reflects the importance of the TF regulon in each cell type. **i,** Heatmap showing the inferred regulon activity (from both scRNA-seq and scRNA-seq data >10 PCW male samples) of the transcription factors (x-axis) involved in the establishment of cellular diversity in the Wolffian duct-derived mesenchymal cells (y-axis). The colour scale is proportional to the expression of the TF while the size of the dot reflects the importance of the TF regulon in each cell type.

**Extended Data Figure 8. Prioritisation of spatially-variable transcription factors and cell-cell communication events along the Müllerian and Wolffian rostro-caudal axes. a,** Schematic representation of the computational workflow used to prioritise spatially-variable genes along the Müllerian rostro-caudal axis from *10x Visium* and *In Situ* Sequencing (*ISS*) female samples >10 PCW. **b,** Smoothed splines of the expression of *LGR5* and *TSPAN8* along the imputed Müllerian rostro-caudal axis in scRNA-seq data modelled with *TradeSeq*^107^. **c,** (left) Venn diagram showing the intersection of genes that are identified as spatially-variable along the measured (in *10x Visium*) and imputed (in scRNA-seq) Müllerian rostro-caudal axes for mesenchymal spots/cells (top) and epithelial spots/cells (bottom). (right) Distribution of cosine similarities between the smoothing splines of the genes that are identified by both data modalities for mesenchymal spots/cells (top) and epithelial spots/cells (bottom). **d,** Heatmap showing the min-max normalised expression measured in scRNA-seq data of prioritised spatially variable epithelial transcription factors (x-axis) along the binned imputed Müllerian rostro-caudal axis (y-axis). **e,** Schematic representation of the computational workflow used to prioritise spatially-variable genes along the Wolffian rostro-caudal axis from *10x Visium* male samples >10 PCW. **f,** Schematic representation of the computational workflow used to prioritise spatially-variable mesenchymal-epithelial interactions along the imputed Müllerian and measured Wolffian rostro-caudal axes from female and male samples >10 PCW. **g,** Heatmap showing the min-max normalised expression measured in scRNA-seq data of epithelial partners of the spatially variable mesenchymal ligand/receptors (x-axis) along the binned imputed Müllerian rostro-caudal axis (y-axis). **h,** Heatmap showing the min-max normalised expression measured in *10x Visium* data of epithelial partners of the spatially variable mesenchymal ligand/receptors (x-axis) along the binned measured Wolffian rostro-caudal axis (y-axis). **i,** Batch corrected, t-distributed Stochastic Neighbour Embedding (t-SNE) embedding of downsampled Müllerian-derived epithelial cells (n = 2345) from female samples >10 PCW profiled with scATAC-seq coloured by stage (measured in PCW) and cell type. **j,** Heatmap showing the inferred regulon activity (from both scRNA-seq and scRNA-seq data >10 PCW female samples) of the transcription factors (x-axis) involved in the establishment of cellular diversity in the Müllerian duct-derived epithelial cells (y-axis). The colour scale is proportional to the expression of the TF while the size of the dot reflects the importance of the TF regulon in each cell type. **k,** Batch corrected, t-SNE embedding of Wolffian-derived epithelial cells (n = 1213) from male samples >10 PCW profiled with scATAC-seq coloured by stage (measured in PCW) and cell type. **l,** Heatmap showing the inferred regulon activity (from both scRNA-seq and scRNA-seq data >10 PCW male samples) of the transcription factors (x-axis) involved in the establishment of cellular diversity in the Wolffian duct-derived epithelial cells (y-axis). The colour scale is proportional to the expression of the TF while the size of the dot reflects the importance of the TF regulon in each cell type.

**Extended Data Figure 9. Intra-organ regionalisation of the fallopian tube and epididymis. a,** Schematic representation of the computational workflow used to prioritise spatially-variable genes along the fallopian tube axis in *10x Visium* female samples >10 PCW. **b,** (top) Haematoxylin and eosin (H&E) stained images of the three capture areas of a 37 years old fallopian tube sample highlighted in **Figure 6e** profiled with *10x Visium*. (middle) H&E stained images of the three capture areas with spots coloured by histological annotation. (bottom) Normalised, log-transformed expression of *MUC6* in each *10x Visium* spot of the three capture areas. **c,** Dot plot showing the variance-scaled, log-transformed expression of genes (x-axis) whose expression increases rostro-caudally along the fetal fallopian tube in the epithelial spots of the three capture areas of the adult fallopian tube (y-axis). **d,** Schematic representation of the computational workflow used to prioritise spatially-variable genes along the epididymal axis in *10x Visium* male samples > 10 PCW. **e,** (left) H&E stained image of a representative section of a 16 PCW epididymis and vas deferens sample profiled with *10x Visium* along side spatial mapping of ciliated cells from the scRNA-seq dataset onto the corresponding *10x Visium* slide using *cell2location*^104^. (right) H&E stained image of a representative section of a 14 PCW epididymis and vas deferens sample profiled with *10x Visium* along side spatial mapping of ciliated cells from the scRNA-seq dataset onto the corresponding *10x Visium* slide using *cell2location*^104^. Estimated cell type abundance (colour intensity) in each Visium spot is shown over the H&E image.

## Data and code availability

We provide user-friendly access to our scRNA-seq resource at www.reproductivecellatlas.org. All the raw and processed sequencing data generated in this study are currently being deposited to ArrayExpress (scRNA-seq, scATAC-seq, *10x Visium*) and BioImageArchive (*ISS*, RNAscope). The code used to perform the analyses presented in the manuscript can be found at https://github.com/ventolab/dev-rep-tract/.

## Acknowledgements

This publication is part of the Human Cell Atlas – www.humancellatlas.org/publications/. We are thankful to the Sanger Cellular Generation and Phenotyping (CGaP) Core Facility and the Sanger Core Sequencing pipeline for support with sample processing and sequencing library preparation. We thank Antonio García (https://www.bio-graphics.es/) for his invaluable help with conceptualising and making the illustrations that are part of this manuscript; Kui Hua for his advice on statistical testing; Aidan Maartens for proofreading and providing advice on the narrative of the manuscript. We are also thankful to the IBSA Foundation for scientific research for supporting V.L. and E.R.R-M. with their yearly fellowship for early career scientists.

## Author information

V.L, L.G-A, J.C.M and R.V-T conceived and designed the experiments and analyses. V.L analysed the data with contributions from N.Y, T.L, and M-A.J. C.I.M and C.S-S performed sample processing. C.I.M, E.T, and K.R performed the imaging experiments. M.M derived the organoids and E.R.R-M, I.K conducted the perturbation experiment. X.H. and B.C. performed sample dissections. V.L, L.G-A, J.C.M and R.V-T interpreted the data. V.L, L.G-A, J.C.M and R.V-T wrote the manuscript. L.G-A, J.C.M and R.V-T supervised the work. All authors read and approved the manuscript.

## Methods

### Patient samples

All tissue samples used for this study were obtained with written informed consent from all participants in accordance with the guidelines in The Declaration of Helsinki 2000. The human embryonic and fetal material was provided by the Joint MRC / Wellcome Trust (grant# MR/R006237/1 and MR/X008304/1) Human Developmental Biology Resource (HDBR, http://www.hdbr.org), with appropriate maternal written consent and approval from the Fulham Research Ethics Committee (REC reference 18/LO/0822 and 23-LO/0312) and Newcastle & North Tyneside 1 Research Ethics Committee (REC reference 18/NE/0290). The HDBR is regulated by the UK Human Tissue Authority (HTA; www.hta.gov.uk) and operates in accordance with the relevant HTA Codes of Practice. This research was also supported by the NIHR Cambridge Biomedical Research Centre (NIHR203312). The views expressed are those of the authors and not necessarily those of the NIHR or the Department of Health and Social Care.

### Assignment of developmental stage

Embryos up to eight post-conception weeks (PCW) were staged using the Carnegie staging method^108^. At stages beyond 8 PCW, age was estimated from measurements of foot length and heel-to-knee length and compared with the standard growth chart^109^. A piece of skin, or where this was not possible, chorionic villi tissue was collected from every sample for quantitative PCR analysis using markers for the sex chromosomes and the following autosomes 13, 15, 16, 18, 21, 22, which are the most commonly seen chromosomal abnormalities. All samples were karyotypically normal.

### Tissue processing

All tissues for sequencing and spatial work were collected in HypoThermosol**®**FRS Preservation solution (Sigma-Aldrich) and stored at 4°C until processing. Tissue dissociation was conducted within 24 hours of tissue retrieval with the exception of tissues that were cryopreserved and stored at -80°C (see **Supplementary Table 1**).

We used the previous protocol optimised for gonadal dissociation and available at protocols.io^110^. In short, tissues were cut into <1 mm^3^ segments before being digested with a mix of Trypsin-EDTA 0.25% and DNaseI (0.1mg/ml) for 5-15 minutes at 37°C with intermittent shaking. Samples > 17 PCW were digested using a combination of collagenase and Trypsin/EDTA, a protocol adapted from Wagner et al.,^110,111^. In short, samples were first digested with a mix of Collagenase 1A (1 mg/ml), DNaseI (0.1 mg/ml) and Liberase TM (50 µg/ml) for 45 minutes at 37°C with rotation. Cell solution was further digested with Trypsin/EDTA 0.25% for 10 minutes at 37°C with rotation. In both protocols, digested tissue was passed through a 100 µm filter, and cells collected by centrifugation (500 x g for 5 minutes at 4°C). Cells were washed and resuspended in PBS-BSA 0.04% prior to cell counting.

### Single nuclei suspension

Single-nuclei suspensions were isolated from dissociated cells when performing scATAC-seq, following the manufacturers’ instructions, and from frozen tissue sections when performing multiomic snRNA-seq/scATAC-seq. For the latter, thick (300 µm) sections were cryosectioned, and kept in a tube on dry ice until subsequent processing. Nuclei were released via Dounce homogenisation as described in detail in the protocols.io.

### Tissue cryopreservation

Fresh tissue was cut into <1 mm^3^ segments before being resuspended with 1 ml of ice cold Cryostor solution (CS10) (C2874-Sigma). The tissue was frozen at -80°C decreasing the temperature approximately 1°C per minute. Detailed protocol available at https://www.protocols.io/view/tissue-freezing-in-cryostor-solution-processing-bgsnjwde.

### Tissue freezing

Fresh tissue samples of human developing reproductive tract were embedded in cold OCT medium and flash frozen using a dry ice-isopentane slurry.

### Haematoxylin and Eosin (H&E) staining and imaging

Fresh frozen sections were removed from -80°C storage and air dried before being fixed in 10% neutral buffered formalin for 5 minutes. After rinsing with deionised water, slides were dipped in Mayer’s Haematoxylin solution (QPath) for 90 seconds. Slides were completely rinsed in 4-5 washes of deionised water, which also served to blue the haematoxylin. Aqueous eosin 1% (Leica) was manually applied onto sections with a pipette and rinsed with deionised water after 1-3 seconds. Slides were dehydrated through an ethanol series (70%, 70%, 100%, 100%) and cleared twice in 100% xylene. Slides were coverslipped and allowed to air dry before being imaged on a Hamamatsu Nanozoomer 2.0HT digital slide scanner.

### Multiplexed smFISH and high-resolution imaging

Large tissue section staining and fluorescent imaging was conducted largely as described previously^112^. Sections were cut from fresh frozen or fixed frozen samples embedded in OCT at a thickness of 10 μm using a cryostat, placed onto SuperFrost Plus slides (VWR) and stored at -80°C until stained. Tissue sections were then processed using a Leica BOND RX to automate staining with the RNAscope Multiplex Fluorescent Reagent Kit v2 Assay (Advanced Cell Diagnostics, Bio-Techne), according to the manufacturers’ instructions. Probes are found in **Supplementary Table 3**. Prior to staining, human fresh frozen sections were post-fixed in 4% paraformaldehyde in PBS for 15 minutes at 4°C, then dehydrated through a series of 50%, 70%, 100%, and 100% ethanol, for 5 minutes each. Following manual pre-treatment, automated processing included epitope retrieval by protease digestion with Protease IV for 30 minutes prior to probe hybridisation. Subsequently, the automated processing for these sections included heat-induced epitope retrieval at 95°C for 5 minutes in buffer ER2 and digestion with Protease III for 15 minutes prior to probe hybridisation. Tyramide signal amplification with Opal 520, Opal 570, and Opal 650 (Akoya Biosciences) and TSA-biotin (TSA Plus Biotin Kit, Perkin Elmer) and streptavidin-conjugated Atto 425 (Sigma Aldrich) was used to develop RNAscope probe channels.

Stained sections were imaged with a Perkin Elmer Opera Phenix High-Content Screening System, in confocal mode with 1 μm z-step size, using a 20X (NA 0.16, 0.299 μm/pixel); 40X (NA 1.1, 0.149 μm/pixel); or 63X (NA 1.15, 0.091 μm/pixel) water-immersion objective. Channels: DAPI (excitation 375 nm, emission 435-480 nm), Atto 425 (ex. 425 nm, em. 463-501 nm), Opal 520 (ex. 488 nm, em. 500-550 nm), Opal 570 (ex. 561 nm, em. 570-630 nm), Opal 650 (ex. 640 nm, em. 650-760 nm).

### Image stitching

Confocal image stacks were stitched as two-dimensional maximum intensity projections using proprietary Acapella scripts provided by Perkin Elmer.

### 10x Genomics Chromium GEX library preparation and sequencing

For the scRNA-seq experiments, cells were loaded according to the manufacturer’s protocol for the Chromium Next GEM Single Cell 5′ v2 (DUAL) Kit (10X Genomics) to attain between 2,000 and 10,000 cells per reaction. Library preparation was carried out according to the manufacturer’s protocol. Libraries were sequenced, aiming at a minimum coverage of 20,000 raw reads per cell, on the Illumina HiSeq 4000 or Novaseq 6000 systems; using the sequencing format; read 1: 26 cycles; i7 index: 8 cycles, i5 index: 0 cycles; read 2: 98 cycles.

For the scATAC-seq and multimodal snRNA-seq/scATAC-seq experiments, cells were loaded according to the manufacturer’s protocol for the Chromium Single Cell ATAC v2 and Chromium Next GEM Single Cell Multiome ATAC+Gene Expression (10X Genomics) to attain between 2,000 and 10,000 cells per well. Library preparation was carried out according to the manufacturer’s protocol. Libraries for scATAC-seq were sequenced on Illumina NovaSeq 6000, aiming at a minimum coverage of 10,000 fragments per cell, with the following sequencing format; read 1: 50 cycles; i7 index: 8 cycles, i5 index: 16 cycles; read 2: 50 cycles.

### 10x Genomics Visium library preparation and sequencing

10 µm cryosections were cut and placed on Visium slides. These were processed according to the manufacturer’s instructions. Briefly, sections were fixed with cold methanol, stained with haematoxylin and eosin and imaged on a Hamamatsu NanoZoomer 2.0HT before permeabilization (24-30 mins), reverse transcription and cDNA synthesis using a template-switching protocol. Second-strand cDNA was liberated from the slide and single-indexed libraries prepared using a 10x Genomics PCR-based protocol. Libraries were sequenced (1 per lane on a HiSeq4000), aiming for 300M raw reads per sample, with the following sequencing format; read 1: 28 cycles, i7 index: 8 cycles, i5 index: 0 cycles and read 2: 91 cycles.

### 10x Genomics Visium CytAssist library preparation and sequencing

10 µm cryosections were collected onto SuperFrost Plus slides (VWR) and processed according to the 10x CytAssist protocol (CG000614 and CG000495). Briefly, sections were methanol fixed, H&E stained and imaged on a Hamamatsu Nanozoomer 2.0HT. After destaining, human whole transcriptome probe pairs were hybridised and ligated to the tissue RNA. The ligation products were then released and captured onto Visium slides, using a CytAssist instrument. The probes were then extended to incorporate the spatial barcodes from the Visium slide, eluted and prepared into a dual-indexed library. Libraries were sequenced (4 samples per Illumina Novaseq SP flow cell) aiming for a minimum 25,000 read pairs per spot, with the sequencing format; read 1: 28 cycles, i7 index: 10 cycles, i5 index: 10 cycles and read 2S: 90 cycles.

### Customised In situ sequencing pipeline

In situ sequencing was performed using the 10X Genomics CARTANA HS Library Preparation Kit (1110-02, following protocol D025) and In Situ Sequencing Kit (3110-02, following protocol D100), which comprise a commercialised version of HybISS^113^. A reproductive tract section was fixed in 3.7% formaldehyde (Merck 252549) in PBS for 30 minutes, washed twice in PBS for 1 minute each, permeabilized in 0.1 M HCl (Fisher 10325710) for 5 minutes, and washed twice again in PBS, all at room temperature. Following dehydration in 70% and 100% ethanol for 2 minutes each, a 50, 100, or 150 μl volume (depending on the size of the section) SecureSeal hybridisation chamber (Grace Bio-Labs GBL621505-20EA) was adhered to the slide and used to hold subsequent reaction mixtures. Following rehydration in buffer WB3, probe hybridisation in buffer RM1 was conducted for 16 hours at 37°C. The 171-plex probe panel included 5 padlock probes per gene, the sequences of which are proprietary (10X Genomics CARTANA). The section was washed with PBS-T (PBS with 0.05% Tween-20) twice, then with buffer WB4 for 30 minutes at 37°C, and thrice again with PBS-T. Probe ligation in RM2 was conducted for 2 hours at 37°C and the section washed thrice with PBS-T, then rolling circle amplification in RM3 was conducted for 18 hours at 30°C. Following PBS-T washes, all rolling circle products (RCPs) were hybridised with LM (Cy5 labelling mix with DAPI) for 30 minutes at room temperature, the section was washed with PBS-T and dehydrated with 70% and 100% ethanol. The hybridisation chamber was removed and the slide mounted with SlowFade Gold Antifade Mountant (Thermo S36937).

Imaging of Cy5-labelled RCPs at this stage acted as a QC step to confirm RCP (’anchor’) generation and served to identify spots during decoding. Imaging was conducted using a Perkin Elmer Opera Phenix Plus High-Content Screening System in confocal mode with 1 μm z-step size, using a 63X (NA 1.15, 0.097 μm/pixel) water-immersion objective and 7% overlap between adjacent tiles. Channels: DAPI (excitation 375 nm, emission 435-480 nm), Atto 425 (ex. 425 nm, em. 463-501 nm), Alexa Fluor 488 (ex. 488 nm, em. 500-550 nm), Cy3 (ex. 561 nm, em. 570-630 nm), Cy5 (ex. 640 nm, em. 650-760 nm).

Following imaging, the slide was de-coverslipped vertically in PBS (gently, with minimal agitation such that the coverslip ‘fell’ off to prevent damage to the tissue). The section was dehydrated with 70% and 100% ethanol, and a new hybridisation chamber secured to the slide. The previous cycle was stripped using 100% formamide (Thermo AM9342), which was applied fresh each minute for 5 minutes, then washed with PBS-T. Barcode labelling was conducted using two rounds of hybridisation, first an adapter probe pool (AP mixes AP1-AP6, in subsequent cycles), then a sequencing pool (SP mix with DAPI, customised with Atto 425 in place of Alexa Fluor 750), each for 1 hour at 37°C with PBS-T washes in between and after. The section was dehydrated, the chamber removed, and the slide mounted and imaged as previously. This was repeated another five times to generate the full dataset of 7 cycles (anchor and 6 barcode bits).

### Derivation of fetal uterine organoids and perturbations with doxycycline

Fetal uterine organoids were derived from an 18 PCW fetal reproductive tract sample (developing uterus, cervix, and vagina) following tissue dissociation as described above. The single-cell suspension was washed in Advanced DMEM/F12 (Gibco, 12634-010), centrifuged and the cell pellet resuspended in Matrigel (Corning, 356231) at around 1:3 ratio (pellet volume : Matrigel volume). The organoids were cultured as described previously^114^, forming in 25 µL Matrigel domes in 48-well plates covered by 250 µL of basal endometrial organoid media (Advanced DMEM/F12 (Gibco, 12634-010), 1% GLUTAMAX (Gibco, 35050061), 1% Insulin-Transferrin-Selenium (ITS -G)(Gibco, 41400045), 100 µg/mL Primocin (Invivogen, ant-pm-1), 1X B27-Vitamin A (Life Technologies, 12587010), 1x N2 (Life Technologies, 17502048), 1.25 mM N-Acetyl-L-cysteine (Sigma-Aldrich, A9165-5G), 2 mM Nicotinamide (Sigma, N0636-100G), 2 ng/mL Recombinant Human FGF-basic (154 a.a.) (Peprotech, 100-18B), 500 ng/mL recombinant human R-spondin-1 (R&D Systems, 4645-RS-01M/CF), 10 µM SB202190 (p38i) (StemCell Technologies, 72632), 500nM A83-01 (Tocris, 2939), 50 ng/mL recombinant human EGF (Peprotech, AF-100-15), 10 ng/mL recombinant human FGF-10 (Peprotech, 100-26) and 100 ng/mL recombinant human Noggin (Peprotech, 120-10C))^114^. The media was supplemented with 10 µM of ROCK inhibitor Y-27632 (Millipore, 688000) for the first 2 days of organoid line establishment and changed every 2-3 days. Once the majority of organoids reached 30-60 µm in diameter, the media was removed and replaced with 250 µL of ice-cold Cell Recovery Solution (Corning, CLS354253) and incubated for 1 h at 4°C. Contents of the wells were then collected, centrifuged, supernatant discarded and organoids resuspended and frozen as whole intact organoids in Cryostor solution CS10 at -80°C, and later transferred and stored in liquid nitrogen.

The frozen organoids were thawed in a water bath at 37 °C. Thawed organoids were transferred to a 50ml Falcon tube and 5 mL of DMEM/F12 was added dropwise The suspension was centrifuged at 800g for 2 min at 4°C. The pellet was resuspended in 100 µL of ice-cold Matrigel and expanded in 30 µL Matrigel (Corning, CLS356255) domes cultured in 6-well plates with basal endometrial organoid media without the addition of 17-β estradiol for two passages at 37 °C (21% O_2_, 5% CO_2_). After each passage, the media was supplemented with 10 µM of ROCK inhibitor Y-27632 (Millipore, 688000) for two days. The media was changed every 2-3 days for fresh basal media.

The organoids were split and passaged approximately every 5-7 days, depending on their size and density. 2 mL of TrypLE™ Express Enzyme (Gibco, 12604013) were added to each well, and a cell scraper was used to detach the domes from the bottom of the well. The organoid suspension was collected with a non-Wide bore 1000 µL tip and transferred to a 15 mL Falcon tube. The organoids were dissociated into cell clumps by pipetting the solution up and down 15 times using a 1000 µL tip, followed by incubation at 37 °C for 6 min. 2 mL of Advanced DMEM/F12 were added and the organoid suspension was pipetted up/down 10 more times with the 1000 µL tip. The suspension was centrifuged at 800*g* for 2 min at 4°C. The supernatant was aspirated as close to the pellet as possible. 30 µL of cold Matrigel per desired dome were added and the pellet was slowly resuspended to evenly distribute the cells while avoiding bubbles. To form the domes, 30 µL domes were pipetted out per well of a 6 well plate. The domes were placed in the incubator for 10 min at 37 °C, followed by the addition of 2 mL of warm media supplemented with ROCK inhibitor per well.

For drug treatment experiments, organoids were passages 48 hours before addition of the drug. Organoids were plated in 30µl domes as described above in two technical replicates. Control wellsreceived basal media. Treated wells received basal media supplemented with 3 µg/mL (5.8 µM) of doxycycline hyclate (Sigma, D5207-1G). This concentration was chosen as it represents drug blood serum levels during doxycycline oral treatment^115,116^. On day 4 of doxycycline treatment, organoids were dissociated into a single-cell suspension. The dissociation process followed the same steps as the passaging. TrypLE digestion step was extended to ∼25 minutes at 37°C. The suspension was visually inspected to ensure that organoids were digested into single cells. After the last centrifugation step, the cell pellet was resuspended in 200µl 1% PBS/BSA and mixed well to ensure full reconstitution. Cell suspension was filtered through a 40µm filter. Live cells were counted using Trypan blue. The cells were loaded into a 10X Genomics Chromium chip as described by the Chromium Next GEM Single Cell 5′ v2 (DUAL) Kit.

### Analysis of scRNA-seq data

#### Per-library analyses

For each sequenced scRNA-seq library, we performed read alignment to the human reference genome (GRCh38 2020-A), mRNA quantification and initial quality control using STARsolo^117^ with default parameters. Due to the scarcity of human cell type markers, each library was first analysed independently before integrating them to have a way of formally evaluating the quality of the integration. Briefly, we used *Scrublet*^118^ for cell doublet calling with a two-step diffusion doublet identification followed by Bonferroni-FDR correction and a significance threshold of 0.01, as described in. Genes expressed by less than three cells were excluded, as were cells expressing fewer than 1500 genes, more than 20% mitochondrial genes, or with more than 40% of *scrublet* score.

After converting the expression space to log(CPM/100 + 1), the *anndata* object was transposed to gene space to identify cell cycling genes in a data-driven manner, as described in^119,120^. PCA, neighbour identification and partition-based *Leiden* clustering^121^ were performed on the gene space, and then the members of the gene cluster including known cycling genes (*CDK1*, *MKI67*, *CCNB2* and *PCNA*) were flagged as the data-derived cell cycling genes, and discarded in the downstream analysis. Back in the cell space, we identified highly variable genes, performed PCA, computed the neighbourhood graph, *Leiden* clustering^121^ and Uniform Manifold Approximation and Projection (*UMAP*)^122^ for visualisation in two dimensions. The per-library computational analysis workflow described so far was wrapped in a Nextflow^123^ pipeline with two processes to enable parallelisation and reproducibility.

To find the genes characteristic of each cluster, we performed term frequency - inverse document frequency (*TF-IDF*), a method borrowed from natural language processing which reflects how important a word (gene) is to a document (cluster) in a corpus (dataset), as implemented in the R library *SoupX*^124^. Annotations were only finalised when analysing spatially-resolved transcriptomics data (both 10x Visium and In Situ Sequencing). A detailed explanation of the cell types identified alongside their marker genes can be found in **Supplementary Note 1**.

#### Integration of scRNA-seq libraries

After having annotated each sample separately and realising the significant differences in cell type composition across samples, we generated three integrated views that best preserve the biological heterogeneity of this system: <= 10 PCW male and female samples together (while the sexually unmatched ducts are still present and the first signs of regionalisation of the ducts appear); > 10 PCW males; > 10 PCW females. The variational autoencoder-based method *scVI*^125^ trained on the 7,500 most highly variable genes and with 30 latent variables and 2 hidden layers was then applied to mitigate batch effects across donors in each of the three views. To evaluate the integrated manifold, we then overlaid the per-sample annotations and confirmed that the biological signal is preserved while correcting for donor-specific effects. Moreover, for a more quantitative evaluation of the integration results, we computed the Shannon entropy per *Leiden* cluster of the per-sample cell type annotations as well as the donor and sex labels, following the approach taken by^126^. Clusters with donor label entropy equal to 0 (i.e. donor-specific clusters) were removed from further analysis. Each cluster was then annotated based on majority voting (>= 40%) of the cell type labels. Clusters showing high cell type label entropy (i.e. < 40% of cells in the cluster having the same cell type label) were further investigated and annotated according to their most expressed *TF-IDF* markers.

Finally, for visualisation purposes only, all libraries were integrated with scVI (7,500 HVGs, 60 latent variables) and cell type labels were overlaid from the per-view annotations. Variations in cell type proportions across developmental time were visualised with an area plot.

#### Per-organ analyses

The cellular and molecular features of the establishment of sexual dimorphism in each organ of the developing human reproductive tract were investigated by performing sub-analyses on the relevant cell types:

- All cell types in the male and female external genitalia during the “masculinisation programming window” (8-14 PCW)
- Coelomic epithelium, Müllerian duct epithelium and mesenchyme during the period of Müllerian duct emergence (6-8 PCW)
- Mesenchymal and epithelial cells from the differentiating Müllerian and Wolffian ducts (>10 PCW), resulting in 4 sub-analyses (Müllerian-derived mesenchyme, Müllerian-derived epithelium, Wolffian-derived mesenchyme, Wolffian-derived epithelium).

In all these per-organ analyses, preprocessing was carried out analogously to what described in the per-library analysis section, and *Harmony*^127^ (*theta* = 0) was employed to correct for batch effects.

#### Human-mouse comparison of the external genitalia

We leveraged the availability of an annotated scRNA-seq dataset of the mouse genital tubercle (comprising two male and female samples for each of three developmental stages: E14.5, E16.6, and E18.5)^36^ to identify the transcriptional regulators underpinning sexual dimorphism in the *corpus spongiosum* in each species. To define a shared feature space, we first took the set of orthologous genes expressed in at least 10 cells in both species. From the set of common orthologous genes, we then computed the top 4000 HVGs in each species and took their intersection (∼2.7 k genes). Using the intersection of HVGs, batch effects due to different donors/mice were corrected by means of Harmony^127^, and a Milo^128^ object was computed on each species’ batch-corrected embedding. Each neighbourhood from a species was matched to its *k* (k = 30) closest neighbourhoods in the other species (in both the mouse-to-human and human-to-mouse directions). The final matches were formed by identifying the mutually nearest pairs of neighbourhoods that appeared in both directions^129^. Each matched neighbourhood was then annotated by majority voting of the labels of the cells in the neighbourhood. Matching across cell type labels was visualised with an alluvial plot. We then considered the unique combination of matching cell type labels as the harmonised cell type annotations across species and focussed on the mouse cells that matched the human *corpus spongiosum* label for differential expression analysis between males and females.

#### Differential expression analysis in the human and mouse genital tubercle

We conducted differential expression analysis between male and female samples within the human and mouse *corpus spongiosum* and human urethral epithelium using *PyDESeq2*^105^. Only samples between 8 and 14 PCW (the so-called “masculinisation programming window”) were included in the analysis. Genes mapping to the Y chromosome were excluded from differential expression testing. Results of the differential expression analysis were visualised with a Volcano plot showing the genes with |logFC > 1.5| and adjusted p-value < 0.05.

#### Cell-cell interaction analysis from scRNA-seq data

Sexually dimorphic cell-cell interactions between the mesenchymal *corpus spongiosum* and the urethral epithelium during the “masculinisation programming window” (8-14 PCW) were inferred with CellphoneDB v5.1^45^ using method 3 (differential expression analysis). After splitting both cell clusters into male and female, differentially expressed genes for each cell type-sex combination were identified using the *FindAllMarkers()* function in *Seurat*^130^, with only positive markers being considered. Only genes expressed in at least 10% of the cells in a cell type-sex combination were considered for this analysis. The search for interaction was restricted to cell types within the same sex (which was passed as input to the method in the “microenvironments” file).

Wolffian-to-Müllerian duct signalling during the developmental time window of Müllerian duct emergence (6-8 PCW) was also explored with CellphoneDB v5.1 using method 3. Differentially expressed genes per cell type (i.e. Wolffian mesenchyme, Wolffian epithelium, Müllerian mesenchyme, Müllerian epithelium) were similarly computed using the *FindAllMarkers()* function in *Seurat*^130^ (with only positive markers being considered), and only genes expressed in at least 10% of the cells in a cell type were considered for this analysis. All cell types belong to the same “microenvironment”.

#### Trajectory inference and differential expression along trajectories

The trajectory inference method *Slingshot*^47^ was applied to recover the lineages originating from the coelomic epithelium during Müllerian duct emergence (6-8 PCW). The pseudotime ordering of the cells along with the weighted assignment of each cell to the three lineages were then used as input for *TradeSeq*^107^ to extract genes that are differentially expressed along a lineage or between lineages with the *associationTest()* function.

#### Inference of clinically-approved drugs potentially disrupting reproductive epithelia

Using *drug2cell*^81^, we derived drug scores for compounds in the CHEMBL database by averaging the expression levels of target genes in each cell on the three views of the dataset independently. We then performed a Wilcoxon rank-sum test to identify significant differences in drug scores between each reproductive epithelium (i.e. Müllerian duct epithelium, Wolffian duct epithelium, urogenital sinus epithelium, and urethral epithelium in <10 PCW female and male samples; fallopian tube epithelium, uterocervix epithelium, upper vagina epithelium, vaginal plate epithelium, and urethral epithelium in >10 PCW female samples; epididymis epithelium, vas deferens epithelium, prostate epithelium, urethral epithelium in >10 PCW males) and all other reproductive-specific cells in the dataset. Results were filtered based on adjusted p-value (< 0.01), log-fold changes (> 2), and rank scores to select the most significant drugs associated with the target cell type. An additional filtering step was performed to exclude drugs whose target genes are not specific to the target cell type and require that the targets are expressed in at least 10% of cells in the target cell type.

#### In vivo - in vitro comparison

Epithelial cells (uterocervix, Müllerian vagina, vaginal plate) from the 18 PCW female sample from which the organoid line was derived were used to train a *CellTypist*^131^ model with the top 200 genes per cell type (as ranked by their absolute regression coefficients associated with each cell type) as features. The trained model then used to predict the cell type labels in the organoid sample (untreated and treated with doxycycline) and the predicted probabilities were visualised using a dot plot.

### Analysis of scATAC-seq data

#### Per-library analyses

scATAC-seq libraries were processed (read filtering, alignment, barcode counting, and cell calling) with 10x Genomics Cell Ranger ATAC pipeline v.2 using the pre-built 10x GRCh38 genome (version 3.1.0) as reference. Similarly to RNA, we analysed each ATAC library independently until cell type annotation in order to evaluate the quality of the subsequent integrations using the *ArchR* framework^132^. Cells with less than 3500 fragments or a minimum transcription start site enrichment of 10 were filtered out, as were cells deemed as doublets. Dimensionality reduction on the *tile matrix* was performed with Latent Semantic Indexing, and the low-dimensional variables were then used to compute the neighbourhood graph, partition-based *Leiden* clustering^121^ and *UMAP* visualisation^122^.

To annotate the cells, we employed Canonical Correlation Analysis to match the *gene activity score matrix* of each scATAC-seq library with the integrated gene expression space of the corresponding view (e.g. if the scATAC-seq library comes from a female sample older than 10 PCW, we use the >10 PCW female integrated scRNA-seq dataset). For optimal reproducibility and parallelisation, the per-sample scATAC-seq analyses were also wrapped in a *Nextflow* script.

#### Integration of scATAC-seq libraries

The individually annotated samples were then integrated by re-computing Latent Semantic Indexing on the concatenated *tile matrices* and using Mutual Nearest Neighbours^129^ to correct for batch effects. Mutual Nearest Neighbours has proven very effective for scATAC-seq (where we do not have as many samples as scRNA-seq) as it allows to specify the order of integration and hence mitigates batch effects in a more biologically informed fashion.

#### Integrative analysis of scRNA-seq and scATAC-seq data

Combining information from transcriptomics and chromatin accessibility data allows for the prioritisation of transcription factors (TFs) that are likely to be active in each cell type, along with the identification of putative regulatory relationships between TFs and target genes. We sought to do this when investigating the regulatory landscape underlying the process of urethral canalisation in males; Müllerian duct emergence from the coelomic epithelium; patterning of the mesenchyme and epithelium during Müllerian and Wolffian duct differentiation.

Using the fragment files and cell annotations obtained through label transfer from the scRNA-seq data, pseudobulks were generated per cell type. Cell-type specific peaks were called using MACS2^133^ as implemented in the *pycistopic* workflow^106^. A set of consensus peaks was derived through iterative overlapping, and the resulting matrix of cells by consensus peaks was used as input to topic modelling with Latent Dirichlet Allocation (LDA). LDA models were selected according to the metrics provided in *pycistopic*. *Harmony*^127^ was used to correct for the donor effect. The manifold was clustered with the *Leiden* algorithm^121^ and differentially accessible regions (DARs) were computed between the *Leiden* clusters. The union of the DARs and the regions contained in topics (obtained by binarising the region-topic probabilities) serve as the set of regions used to find TF binding motifs with *pycistarget*^106^.

To infer enhancer-driven gene regulatory networks, scRNA-seq and scATAC-seq data were eventually combined into a pseudo-multiome dataset using *SCENIC+*^106^. In essence, *SCENIC+* generates metacells containing cells from both data modalities by randomly sampling cells of the same cell type label. Within a 150kb region around each gene, gradient boosting machines and correlation analysis were employed to infer the relationships between enhancers and genes, as well as between transcription factors and genes. This approach allows for the identification of enhancers that are associated with the regulation of specific target genes and the inference of TFs that potentially contribute to the regulation of these genes.

### Analysis of *10x Visium* data

#### Per-library analyses

Visium data consists of FASTQ sequencing files and a bright-field microscopy image stained with haematoxylin and eosin (H&E) per capture area. The Space Ranger (*v2.0.0*) software provided by 10x Genomics was employed to align the barcoded spot pattern to the H&E tissue image and to differentiate tissue from background. The resulting spot-by-transcript abundance matrix was analysed using the package *squidpy*.

#### Estimation of cell type abundances per Visium spot using matched scRNA-seq data

To deconvolve the transcriptional signal coming from each Visium spot into an estimated abundance of each cell type present in the > 10 PCW females and males views of the scRNA-seq dataset, we applied the Bayesian model *cell2location*^104^. Briefly, *cell2location* first estimates reference cell type signatures from the dissociated scRNA-seq data using negative binomial regression. It then decomposes the spatially resolved Visium RNA count matrices into the reference cell type signatures.

#### Annotation of anatomical and histological features

We used the microscopy H&E images to annotate anatomical structures and histological features independently of gene expression. Using the package *TissueTag*^60^, anatomical features were manually labelled, while histological features were inferred using a random forest classifier trained on a few manually labelled points. Estimated cell type abundances per spot in each anatomical structure (or histological feature) were then averaged and an enrichment score of cell type per anatomical structure (or histological feature) was computed.

#### Scoring of the fetal fallopian tube decreasing signature in adult data

To evaluate whether the rostro-caudal decreasing pattern of expression of the 15 genes identified in the fetal fallopian tube was preserved into adulthood, we generated *10x Visium* spatial transcriptomics data from three areas of an adult fallopian tube (fimbriae, ampulla, and isthmus) and scored the epithelial spots in each capture area for the average expression of this gene signature. Briefly, we used the scanpy^134^ function *sc.tl.score_genes()* to compute the score and then performed the Jonckheere’s trend test^135^ with alternative hypothesis “decreasing” (2000 permutations) to quantify the statistical significance of the trend.

### Analysis of *In Situ* Sequencing data

#### Probe selection

To locate cell types and states as they appear and disappear during the development of the reproductive tract (especially during early stages of development which cannot be assayed with spot-based spatial transcriptomics) we designed a panel of 171 genes. The most unique marker genes per cell type were selected with TF-IDF, and the resulting panel was evaluated for completeness using *geneBasis*^136^. To evaluate completeness at the cell level, *geneBasis* checks for preservation of a cell’s neighbourhood in the k-NN graph built with all the HVGs and the k-NN graph constructed with the gene panel by comparing the distance between each cell and these two sets of nearest neighbours. *geneBasis* was applied to evaluate the same gene panel on the three views of the scRNA-seq data (<= 10 PCW males and females, > 10 PCW males, > 10 PCW females) separately. However, as thoracic *HOX* genes are not cited in the literature as relevant to the differentiation of the reproductive tract, these were not included in the original gene panel. We therefore decided to swap 4 of the genes in the original panels (*DPP4, DPP6, CRISP3, DPP10*) for 4 *HOX* genes (*HOXC4, HOXC6, HOXA7, HOXC10*) and run *ISS* on some samples with this resulting panel instead (**Supplementary Table 3**).

#### Preprocessing

ISS data was preprocessed with a computational pipeline implemented by T.L. in *Nextflow*. Briefly, the first step of the pipeline involves integrating the tiles along the Z-axis using maximal projection for each channel and then stitching them together along the remaining X and Y spatial axes. Because the tissue moves slightly between sequencing rounds, image registration is required to correct for spatial misalignment of the fluorescent spots. This registration is achieved using non-linear optical flow to align the small, Gaussian-like spots in the images. Once the images have been stitched and corrected for misalignment, we use the DAPI signal to segment the nuclei with CellPose^137^. For computational efficiency, images are first sliced into smaller 10,000 x 10,000 pixels patches. Even though there is no membrane protein staining, cell segmentation can be obtained by expanding the segmented nuclei by ∼10/15 pixels to mimic the cytoplasm. Finally, spots appearing across registered coding channels are detected and their intensities are extracted. The intensity values for each spot over imaging cycles are then decoded based on the collection of barcodes in the codebook provided by CARTANA with the PoSTcode algorithm^138^. To minimise false positives, PoSTcode inflates the codebook with an additional background barcode. The stacks of image values, representing the intensities across different channels and imaging cycles, are modelled using a matrix-variate Gaussian Mixture Model, assuming correlations between channels and imaging cycles. Decoded spots are ultimately assigned to segmented cells resulting in a gene expression matrix used for downstream analysis.

#### Cell type annotation of ISS data based on matched scRNA-seq data

While the ISS gene panel was designed to maximise cell type recovery, the throughput of the assay is still too limited to confidently assign cell type identities only by examining the measured gene expression. We therefore leveraged the full transcriptome resolution of the scRNA-seq dataset to increase the confidence in cell type assignment using an approach based on kNN graphs implemented in the *iss-patcher* library^139^. Using this approach, we annotated the cells of ISS samples in a per-view fashion (e.g. ISS samples <= 10 PCW were annotated using the <=10 PCW scRNA-seq reference). In early (<= 10 PCW) samples, ISS cells whose anatomical annotation is “gonad” were excluded from the label transfer algorithm because that there are no gonad cells in the scRNA-seq reference data (by design).

#### Annotation of anatomical structures

The experimental workflow of *ISS* currently does not involve the acquisition of an H&E image of the sample. We overcame the limitation of not having an H&E image by generating a “virtual” RGB image from the gene expression profiles of 3 highly expressed markers (one per R, G, B channel). Major anatomical landmarks were therefore annotated on the “virtual” RGB image based on the morphology and the H&E image of the consecutive section. Histological landmarks were not annotated as they necessitate the texture information captured by H&E staining only. By combining the information about the cell type label of each ISS cell and its annotated anatomical location within the tissue architecture, we then computed an enrichment score (z-score) of each cell type in each anatomical location (separately per view of the data). Such z-score enrichment was computed using only the ISS cells with a cell type label fraction above 0.4 (meaning that 40% of the scRNA-seq neighbours are annotated with the same cell type label). The per-view enrichment scores were visualised via a heatmap.

### Computational representation of the Müllerian and Wolffian rostro-caudal axes

Detailed information about the rationale and implementation can be found in **Supplementary Note 2**, **3**, and **4**; here, we provide a summary.

To study the spatial gene expression patterns along the developing female reproductive tract, we constructed the Müllerian rostro-caudal axis by measuring distances from key anatomical landmarks in spatially-resolved transcriptomics data (*10x Visium Cytassist* and *ISS*), as described in the OrganAxis framework^60^. The axis spans from the fallopian fimbriae to the Müllerian vagina-vaginal plate junction, capturing the differentiation of the Müllerian ducts. In cases where the full length of the uterovaginal canal could not be captured in a single section due to the limitations of the *10x Visium Cytassist* platform’s capture area, consecutive tissue sections were computationally stitched together^140^. This stitching involved manually overlapping consecutive sections using the image processing software Fiji (https://imagej.net/plugins/trakem2/) and applying affine transformations to align the sections, which were then concatenated into a single dataset. The resulting Müllerian rostro-caudal axis was normalised and rescaled, allowing for consistent comparison of gene expression patterns across different samples of the female reproductive tract.

To extend our spatial analysis to single-cell resolution, we projected the Müllerian rostro-caudal axis onto scRNA-seq data by leveraging the single-cell resolution provided by *ISS* technology. We restricted our analysis to mesenchymal and epithelial compartments, which are key to understanding the development of the female reproductive tract. Using a modified *k*-nearest neighbours (kNN) approach implemented in the *iss-patcher*^103^ library, each cell in the scRNA-seq dataset was assigned a position along the Müllerian rostro-caudal axis based on its proximity to cells in the *ISS* data. We then employed the *TradeSeq*^107^ framework to model gene expression along the measured (*10x Visium Cytassist*) and imputed (*scRNA-seq*) Müllerian rostro-caudal axis, treating the axis analogously to pseudotime. Genes showing significant changes in expression along the axis were identified using stringent criteria (p-value < 0.001, logFC > 0.5), and we prioritised those with specific expression in mesenchymal or epithelial cells.

In parallel, to investigate gene expression patterns along the male reproductive tract, we constructed the Wolffian rostro-caudal axis, spanning the length of the epididymis and the initial segment of the vas deferens. This axis was derived using data exclusively from *10x Visium Cytassist*, as we lacked sufficient *ISS* male samples for robust imputation of the axis onto scRNA-seq. The axis was similarly defined by calculating the normalised distance between the efferent ductules and the vas deferens. It was then rescaled to maintain consistency with the Müllerian rostro-caudal axis, facilitating comparative analyses. We employed the *TradeSeq*^107^ framework to model gene expression continuously along the Wolffian rostro-caudal axis, and used the same significance thresholds as for the Müllerian rostro-caudal axis to prioritise genes whose expression changes along the differentiating Wolffian ducts.

Prioritised spatially-variable mesenchymal and epithelial genes in the imputed Müllerian rostro-caudal axis and measured Wolffian rostro-caudal axis were then used to filter the transcription factors and cell-cell communication events that likely drive Müllerian and Wolffian regionalisation during fetal development.

## Bibliography

1. Cunha, G. R. et al. Mesenchymal-epithelial interactions in sex differentiation. Hum. Genet. 58, 68–77 (1981).

2. Hyuga, T. et al. Regulatory roles of epithelial-mesenchymal interaction (EMI) during early and androgen dependent external genitalia development. Differentiation 110, 29–35 (2019).

3. Grumbach, M. M. & Ducharme, J. R. The effects of androgens on fetal sexual development: androgen-induced female pseudohermaphrodism. Fertil. Steril. 11, 157–180 (1960).

4. Wilson, J. D., Griffin, J. E. & Russell, D. W. Steroid 5 alpha-reductase 2 deficiency. Endocr. Rev. 14, 577–593 (1993).

5. Cunha, G. R. et al. Development of the human female reproductive tract. Differentiation 103, 46–65 (2018).

6. Robboy, S. J., Kurita, T., Baskin, L. & Cunha, G. R. New insights into human female reproductive tract development. Differentiation 97, 9–22 (2017).

7. Hannema, S. E. & Hughes, I. A. Regulation of Wolffian duct development. Horm. Res. 67, 142–151 (2007).

8. Development of the human prostate. Differentiation 103, 24–45 (2018).

9. Blaschko, S. D., Cunha, G. R. & Baskin, L. S. Molecular mechanisms of external genitalia development. Differentiation 84, 261–268 (2012).

10. Joseph, A., Yao, H. & Hinton, B. T. Development and morphogenesis of the Wolffian/epididymal duct, more twists and turns. Dev. Biol. 325, 6–14 (2009).

11. Du, H. & Taylor, H. S. The Role of Hox Genes in Female Reproductive Tract Development, Adult Function, and Fertility. Cold Spring Harb. Perspect. Med. 6, (2016).

12. Gubbay, J. et al. A gene mapping to the sex-determining region of the mouse Y chromosome is a member of a novel family of embryonically expressed genes. Nature 346, 245–250 (1990).

13. Koopman, P., Gubbay, J., Vivian, N., Goodfellow, P. & Lovell-Badge, R. Male development of chromosomally female mice transgenic for Sry. Nature 351, 117–121 (1991).

14. Arango, N. A., Lovell-Badge, R. & Behringer, R. R. Targeted mutagenesis of the endogenous mouse Mis gene promoter: in vivo definition of genetic pathways of vertebrate sexual development. Cell 99, 409–419 (1999).

15. Shen, W. H., Moore, C. C., Ikeda, Y., Parker, K. L. & Ingraham, H. A. Nuclear receptor steroidogenic factor 1 regulates the müllerian inhibiting substance gene: a link to the sex determination cascade. Cell 77, 651–661 (1994).

16. Behringer, R. R., Finegold, M. J. & Cate, R. L. Müllerian-inhibiting substance function during mammalian sexual development. Cell 79, 415–425 (1994).

17. Cate, R. L. Anti-Müllerian Hormone Signal Transduction involved in Müllerian Duct Regression. Front. Endocrinol. 13, 905324 (2022).

18. Wilson, J. D. The role of 5alpha-reduction in steroid hormone physiology. Reprod. Fertil. Dev. 13, 673–678 (2001).

19. Saravelos, S. H., Cocksedge, K. A. & Li, T.-C. Prevalence and diagnosis of congenital uterine anomalies in women with reproductive failure: a critical appraisal. Hum. Reprod. Update 14, 415–429 (2008).

20. Springer, A., van den Heijkant, M. & Baumann, S. Worldwide prevalence of hypospadias. J. Pediatr. Urol. 12, 152.e1–7 (2016).

21. Guioli, S., Sekido, R. & Lovell-Badge, R. The origin of the Mullerian duct in chick and mouse. Dev. Biol. 302, 389–398 (2007).

22. Orvis, G. D. & Behringer, R. R. Cellular mechanisms of Müllerian duct formation in the mouse. Dev. Biol. 306, 493–504 (2007).

23. Suzuki, K. et al. Embryonic development of mouse external genitalia: insights into a unique mode of organogenesis. Evol. Dev. 4, 133–141 (2002).

24. Schoenwolf, G. C., Bleyl, S. B., Brauer, P. R. & Francis-West, P. H. Larsen’s Human Embryology E-Book. (Elsevier Health Sciences, 2014).

25. Baskin, L. et al. Development of the human penis and clitoris. Differentiation 103, 74–85 (2018).

26. Shen, J. et al. Immunohistochemical expression analysis of the human fetal lower urogenital tract. Differentiation 103, 100–119 (2018).

27. Garcia-Alonso, L. et al. Single-cell roadmap of human gonadal development. Nature 607, 540–547 (2022).

28. Guo, J. et al. Single-cell analysis of the developing human testis reveals somatic niche cell specification and fetal germline stem cell establishment. Cell Stem Cell 28, 764–778.e4 (2021).

29. Li, L. et al. Single-Cell RNA-Seq Analysis Maps Development of Human Germline Cells and Gonadal Niche Interactions. Cell Stem Cell 20, 858–873.e4 (2017).

30. Wamaitha, S. E. et al. Single-cell analysis of the developing human ovary defines distinct insights into ovarian somatic and germline progenitors. Dev Cell 58, 2097–2111.e3 (2023).

31. Taelman, J. et al. Characterization of the human fetal gonad and reproductive tract by single-cell transcriptomics. Dev. Cell 59, 529–544.e5 (2024).

32. Website. 10.1002/jez.1112 doi:10.1002/jez.1112.

33. Dinh, H. Q. et al. Single-cell transcriptomics identifies gene expression networks driving differentiation and tumorigenesis in the human fallopian tube. Cell Rep. 35, 108978 (2021).

34. Carroll, T. J., Park, J. S., Hayashi, S., Majumdar, A. & McMahon, A. P. Wnt9b plays a central role in the regulation of mesenchymal to epithelial transitions underlying organogenesis of the mammalian urogenital system. Dev. Cell 9, (2005).

35. Grote, D., Souabni, A., Busslinger, M. & Bouchard, M. Pax 2/8-regulated Gata 3 expression is necessary for morphogenesis and guidance of the nephric duct in the developing kidney. Development 133, 53–61 (2006).

36. Amato, C. M. & Yao, H. H.-C. Developmental and sexual dimorphic atlas of the prenatal mouse external genitalia at the single-cell level. Proc. Natl. Acad. Sci. U. S. A. 118, (2021).

37. Armfield, B. A. & Cohn, M. J. Single cell transcriptomic analysis of external genitalia reveals complex and sexually dimorphic cell populations in the early genital tubercle. Dev. Biol. 477, 145–154 (2021).

38. Kim, K. S. et al. Expression of the androgen receptor and 5 alpha-reductase type 2 in the developing human fetal penis and urethra. Cell Tissue Res. 307, 145–153 (2002).

39. Wilson, J. D., George, F. W. & Griffin, J. E. The hormonal control of sexual development. Science 211, 1278–1284 (1981).

40. Suzuki, K. et al. Sexually dimorphic expression of Mafb regulates masculinization of the embryonic urethral formation. Proc. Natl. Acad. Sci. U. S. A. 111, 16407–16412 (2014).

41. Sharpe, R. M. Androgens and the masculinization programming window: human-rodent differences. Biochem. Soc. Trans. 48, 1725–1735 (2020).

42. Gene Set - AR. https://maayanlab.cloud/Harmonizome/gene_set/AR/CHEA+Transcription+Factor+Targets.

43. Maurat, E., et al. A novel in vitro tubular model to recapitulate features of distal airways: the bronchioid. bioRxiv 2023.12.06.569771 (2024) doi:10.1101/2023.12.06.569771.

44. Truffi, M. et al. RPTPα controls epithelial adherens junctions, linking E-cadherin engagement to c-Src-mediated phosphorylation of cortactin. J. Cell Sci. 127, 2420–2432 (2014).

45. Troulé, K. et al. CellPhoneDB v5: inferring cell-cell communication from single-cell multiomics data. (2023).

46. Santana Gonzalez, L., et al. Mechanistic Drivers of Müllerian Duct Development and Differentiation Into the Oviduct. Front. Cell Dev. Biol. 9, 605301 (2021).

47. Street, K. et al. Slingshot: cell lineage and pseudotime inference for single-cell transcriptomics. BMC Genomics 19, 1–16 (2018).

48. Tomé, D., Dias, M. S., Correia, J. & Almeida, R. D. Fibroblast growth factor signaling in axons: from development to disease. Cell Commun. Signal. 21, (2023).

49. Le Verche, V. et al. The Somatostatin 2A Receptor Is Enriched in Migrating Neurons during Rat and Human Brain Development and Stimulates Migration and Axonal Outgrowth. PLoS One 4, (2009).

50. Cowell, J. K. LGI1: from zebrafish to human epilepsy. Prog. Brain Res. 213, (2014).

51. Uesaka, T., Nagashimada, M. & Enomoto, H. GDNF Signaling Levels Control Migration and Neuronal Differentiation of Enteric Ganglion Precursors. J. Neurosci. 33, 16372 (2013).

52. Katic, J. et al. Interaction of the cell adhesion molecule CHL1 with vitronectin, integrins, and the plasminogen activator inhibitor-2 promotes CHL1-induced neurite outgrowth and neuronal migration. J. Neurosci. 34, (2014).

53. Zhang, D., Zhou, S. & Liu, B. Identification and Validation of an Individualized EMT-Related Prognostic Risk Score Formula in Gastric Adenocarcinoma Patients. Biomed Res. Int. 2020, (2020).

54. Shi, Q. et al. Mechanisms of Action of Autophagy Modulators Dissected by Quantitative Systems Pharmacology Analysis. Int. J. Mol. Sci. 21, (2020).

55. Zhao, L. et al. COX7A1 suppresses the viability of human non-small cell lung cancer cells via regulating autophagy. Cancer Med. 8, 7762–7773 (2019).

56. Martinelli, I. et al. Galanin promotes autophagy and alleviates apoptosis in the hypertrophied heart through FoxO1 pathway. Redox Biology 40, (2021).

57. Li, N. et al. Targeting ANXA7/LAMP5-mTOR axis attenuates spinal cord injury by inhibiting neuronal apoptosis via enhancing autophagy in mice. Cell Death Discovery 9, 1–16 (2023).

58. Mullen, R. D., Wang, Y., Liu, B., Moore, E. L. & Behringer, R. R. Osterix functions downstream of anti-Müllerian hormone signaling to regulate Müllerian duct regression. Proc. Natl. Acad. Sci. U. S. A. 115, (2018).

59. Park, J. H. et al. Induction of WNT inhibitory factor 1 expression by Müllerian inhibiting substance/AntiMullerian hormone in the Müllerian duct mesenchyme is linked to Müllerian duct regression. Dev. Biol. 386, 227 (2014).

60. Yayon, N. et al. A spatial human thymus cell atlas mapped to a continuous tissue axis. bioRxiv 2023.10.25.562925 (2023) doi:10.1101/2023.10.25.562925.

61. Naylor, R. W. et al. BMP and retinoic acid regulate anterior–posterior patterning of the non-axial mesoderm across the dorsal–ventral axis. Nat. Commun. 7, (2016).

62. Lu, M.-F. et al. prx-1 functions cooperatively with another paired-related homeobox gene,prx-2, to maintain cell fates within the craniofacial mesenchyme. Development 126, 495–504 (1999).

63. Lu, M. F. et al. Paired-related homeobox genes cooperate in handplate and hindlimb zeugopod morphogenesis. Dev. Biol. 205, 145–157 (1999).

64. Capellini, T. D., Zappavigna, V. & Selleri, L. Pbx Homeodomain Proteins: TALEnted regulators of Limb Patterning and Outgrowth. Dev. Dyn. 240, 1063 (2011).

65. Bell, C. C. et al. The Evx1/Evx1as gene locus regulates anterior-posterior patterning during gastrulation. Sci. Rep. 6, 1–11 (2016).

66. Yamagishi, T., Ozawa, M., Ohtsuka, C., Ohyama-Goto, R. & Kondo, T. Evx2-Hoxd13 Intergenic Region Restricts Enhancer Association to Hoxd13 Promoter. PLoS One 2, (2007).

67. St-Jean, G. et al. Targeted ablation of Wnt4 and Wnt5a in Müllerian duct mesenchyme impedes endometrial gland development and causes partial Müllerian agenesis†. Biol. Reprod. 100, 49–60 (2018).

68. Chumduri, C. et al. Opposing Wnt signals regulate cervical squamocolumnar homeostasis and emergence of metaplasia. Nat. Cell Biol. 23, 184 (2021).

69. Laronda, M. M. et al. Diethylstilbestrol induces vaginal adenosis by disrupting SMAD/RUNX1-mediated cell fate decision in the Müllerian duct epithelium. Dev. Biol. 381, 5 (2013).

70. Sullivan, R., Légaré, C., Lamontagne-Proulx, J., Breton, S. & Soulet, D. Revisiting structure/functions of the human epididymis. Andrology 7, 748–757 (2019).

71. Liu, Z. & Li, B. Spatiotemporal expression profile of a putative β propeller WDR72 in laying hens. Mol. Biol. Rep. 40, 5247–5253 (2013).

72. Keating, N. & Quinlan, L. R. Small conductance potassium channels drive ATP-activated chloride secretion in the oviduct. Am. J. Physiol. Cell Physiol. 302, C100–9 (2012).

73. Ulrich, N. D. et al. Cellular heterogeneity of human fallopian tubes in normal and hydrosalpinx disease states identified using scRNA-seq. Dev. Cell 57, 914–929.e7 (2022).

74. Lengyel, E. et al. A molecular atlas of the human postmenopausal fallopian tube and ovary from single-cell RNA and ATAC sequencing. Cell Rep. 41, 111838 (2022).

75. Browne, J. A., Leir, S.-H., Yin, S. & Harris, A. Transcriptional networks in the human epididymis. Andrology 7, 741–747 (2019).

76. Browne, J. A., Yang, R., Leir, S.-H., Eggener, S. E. & Harris, A. Expression profiles of human epididymis epithelial cells reveal the functional diversity of caput, corpus and cauda regions. Mol. Hum. Reprod. 22, 69–82 (2016).

77. Liang, A.-J. et al. The expression of the new epididymal luminal protein of PDZ domain containing 1 is decreased in asthenozoospermia. Asian J. Androl. 20, 154–159 (2018).

78. Li, J. et al. Systematic mapping and functional analysis of a family of human epididymal secretory sperm-located proteins. Mol. Cell. Proteomics 9, 2517–2528 (2010).

79. Reed, C. E. & Fenton, S. E. Exposure to diethylstilbestrol during sensitive life stages: a legacy of heritable health effects. Birth Defects Res. C Embryo Today 99, 134–146 (2013).

80. Hoover, R. N. et al. Adverse health outcomes in women exposed in utero to diethylstilbestrol. N. Engl. J. Med. 365, 1304–1314 (2011).

81. Kanemaru, K. et al. Spatially resolved multiomics of human cardiac niches. Nature 619, 801–810 (2023).

82. Anatomical Therapeutic Chemical (ATC) Classification. https://www.who.int/tools/atc-ddd-toolkit/atc-classification.

83. Lee, H.-R. et al. Molecular mechanism(s) of endocrine-disrupting chemicals and their potent oestrogenicity in diverse cells and tissues that express oestrogen receptors. J. Cell. Mol. Med. 17, 1–11 (2013).

84. Zhao, Y. et al. Phthalate-induced testosterone/androgen receptor pathway disorder on spermatogenesis and antagonism of lycopene. J. Hazard. Mater. 439, 129689 (2022).

85. Fisher, J. S. Environmental anti-androgens and male reproductive health: focus on phthalates and testicular dysgenesis syndrome. Reproduction 127, 305–315 (2004).

86. Evans, N. et al. In vitro activity of a panel of per- and polyfluoroalkyl substances (PFAS), fatty acids, and pharmaceuticals in peroxisome proliferator-activated receptor (PPAR) alpha, PPAR gamma, and estrogen receptor assays. Toxicol. Appl. Pharmacol. 449, 116136 (2022).

87. Abbas, T. O., Mahdi, E., Hasan, A., AlAnsari, A. & Pennisi, C. P. Current Status of Tissue Engineering in the Management of Severe Hypospadias. Front Pediatr 5, 283 (2017).

88. Chan, Y. Y. et al. The current state of tissue engineering in the management of hypospadias. Nat. Rev. Urol. 17, 162–175 (2020).

89. Dyche, W. J. A comparative study of the differentiation and involution of the mullerian duct and wolffian duct in the male and female fetal mouse. J. Morphol. 162, 175–209 (1979).

90. Marečková, M. et al. An integrated single-cell reference atlas of the human endometrium. bioRxiv 2023.11.03.564728 (2023) doi:10.1101/2023.11.03.564728.

91. Chumduri, C. & Turco, M. Y. Organoids of the female reproductive tract. J. Mol. Med. 99, 531–553 (2021).

92. Cyr, D. G. & Pinel, L. Emerging organoid models to study the epididymis in male reproductive toxicology. Reprod. Toxicol. 112, 88–99 (2022).

93. Harnik, Y. et al. A spatial expression atlas of the adult human proximal small intestine. Nature (2024) doi:10.1038/s41586-024-07793-3.

94. Ben-Moshe, S. et al. Spatial sorting enables comprehensive characterization of liver zonation. Nat Metab 1, 899–911 (2019).

95. Hildebrandt, F. et al. Spatial Transcriptomics to define transcriptional patterns of zonation and structural components in the mouse liver. Nat. Commun. 12, 7046 (2021).

96. Maimets, M. et al. Mesenchymal-epithelial crosstalk shapes intestinal regionalisation via Wnt and Shh signalling. Nat. Commun. 13, 715 (2022).

97. Hibbs, K. et al. Differential Gene Expression in Ovarian Carcinoma : Identification of Potential Biomarkers. Am. J. Pathol. 165, 397 (2004).

98. Shen, W. et al. GATA6: a new predictor for prognosis in ovarian cancer. Hum. Pathol. 86, (2019).

99. Taube, E. T. et al. Wilms tumor protein 1 (WT1)--not only a diagnostic but also a prognostic marker in high-grade serous ovarian carcinoma. Gynecol. Oncol. 140, (2016).

100. Erickson, B. K., Conner, M. G. & Landen, C. N. The role of the fallopian tube in the origin of ovarian cancer. Am. J. Obstet. Gynecol. 209, 409–414 (2013).

101. Ogo, F. M. et al. Bisphenol A Exposure Impairs Epididymal Development during the Peripubertal Period of Rats: Inflammatory Profile and Tissue Changes. Basic Clin. Pharmacol. Toxicol. 122, 262–270 (2018).

102. Mechanisms of action of phthalate esters, individually and in combination, to induce abnormal reproductive development in male laboratory rats. Environ. Res. 108, 168–176 (2008).

103. GitHub - Teichlab/iss_patcher: Approximate missing features from higher dimensionality data neighbours. GitHub https://github.com/Teichlab/iss_patcher.

104. Kleshchevnikov, V. et al. Cell2location maps fine-grained cell types in spatial transcriptomics. Nat. Biotechnol. 40, 661–671 (2022).

105. Love, M. I., Huber, W. & Anders, S. Moderated estimation of fold change and dispersion for RNA-seq data with DESeq2. Genome Biol. 15, 1–21 (2014).

106. Bravo González-Blas, C., et al. SCENIC+: single-cell multiomic inference of enhancers and gene regulatory networks. Nat. Methods 20, 1355–1367 (2023).

107. Van den Berge, K., et al. Trajectory-based differential expression analysis for single-cell sequencing data. Nat. Commun. 11, 1–13 (2020).

108. Harper, J. Human Embryology and Teratology. Second Edition. By Ronan O’Rahilly and Fabiola Muller. Ann. Hum. Genet. 60, 533–533 (1996).

109. Hern, W. M. Correlation of fetal age and measurements between 10 and 26 weeks of gestation. Obstet. Gynecol. 63, 26–32 (1984).

110. Sancho, C., Hoo, R. & Vento-Tormo, R. Human embryonic gonad dissociation with Collagenase & Trypsin v3. protocols.io ZappyLab, Inc. 10.17504/protocols.io.bwcipaue (2021).

111. Wagner, M. et al. Single-cell analysis of human ovarian cortex identifies distinct cell populations but no oogonial stem cells. Nat. Commun. 11, 1–15 (2020).

112. Bayraktar, O. A. et al. Astrocyte layers in the mammalian cerebral cortex revealed by a single-cell in situ transcriptomic map. Nat. Neurosci. (2020) doi:10.1038/s41593-020-0602-1.

113. Hilscher, M. M. et al. Spatial and temporal heterogeneity in the lineage progression of fine oligodendrocyte subtypes. BMC Biol. 20, 122 (2022).

114. Boretto, M. et al. Development of organoids from mouse and human endometrium showing endometrial epithelium physiology and long-term expandability. Development 144, 1775–1786 (2017).

115. Newton, P. N. et al. Pharmacokinetics of oral doxycycline during combination treatment of severe falciparum malaria. Antimicrob. Agents Chemother. 49, 1622–1625 (2005).

116. van Roeden, S. E. et al. The effect of measuring serum doxycycline concentrations on clinical outcomes during treatment of chronic Q fever. J. Antimicrob. Chemother. 73, 1068–1076 (2018).

117. Kaminow, B., Yunusov, D. & Dobin, A. STARsolo: accurate, fast and versatile mapping/quantification of single-cell and single-nucleus RNA-seq data. bioRxiv 2021.05.05.442755 (2021) doi:10.1101/2021.05.05.442755.

118. Wolock, S. L., Lopez, R. & Klein, A. M. Scrublet: Computational Identification of Cell Doublets in Single-Cell Transcriptomic Data. Cell systems 8, (2019).

119. Popescu, D.-M. et al. Decoding human fetal liver haematopoiesis. Nature 574, 365–371 (2019).

120. Park, J. E. et al. A cell atlas of human thymic development defines T cell repertoire formation. Science 367, (2020).

121. Traag, V. A., Waltman, L. & van Eck, N. J. From Louvain to Leiden: guaranteeing well-connected communities. Sci. Rep. 9, 1–12 (2019).

122. McInnes, L., Healy, J. & Melville, J. UMAP: Uniform Manifold Approximation and Projection for Dimension Reduction. (2018).

123. Di Tommaso, P. et al. Nextflow enables reproducible computational workflows. Nat. Biotechnol. 35, 316–319 (2017).

124. Young, M. D. & Behjati, S. SoupX removes ambient RNA contamination from droplet-based single-cell RNA sequencing data. Gigascience 9, giaa151 (2020).

125. Gayoso, A., Lopez, R., Xing, G., Boyeau, P. & Wu, K. scvi-tools: a library for deep probabilistic analysis of single-cell omics data. bioRxiv (2021).

126. Sikkema, L. et al. An integrated cell atlas of the lung in health and disease. Nat. Med. 29, 1563–1577 (2023).

127. Korsunsky, I. et al. Fast, sensitive and accurate integration of single-cell data with Harmony. Nat. Methods 16, 1289–1296 (2019).

128. Dann, E., Henderson, N. C., Teichmann, S. A., Morgan, M. D. & Marioni, J. C. Differential abundance testing on single-cell data using k-nearest neighbor graphs. Nat. Biotechnol. 40, 245–253 (2021).

129. Haghverdi, L., Lun, A. T. L., Morgan, M. D. & Marioni, J. C. Batch effects in single-cell RNA-sequencing data are corrected by matching mutual nearest neighbors. Nat. Biotechnol. 36, 421–427 (2018).

130. Hao, Y. et al. Integrated analysis of multimodal single-cell data. Cell (2021) doi:10.1016/j.cell.2021.04.048.

131. Xu, C. et al. Automatic cell-type harmonization and integration across Human Cell Atlas datasets. Cell 186, 5876–5891.e20 (2023).

132. Granja, J. M. et al. ArchR is a scalable software package for integrative single-cell chromatin accessibility analysis. Nat. Genet. 53, 403–411 (2021).

133. Gaspar, J. M. Improved peak-calling with MACS2. bioRxiv 496521 (2018) doi:10.1101/496521.

134. Wolf, F. A., Angerer, P. & Theis, F. J. SCANPY: large-scale single-cell gene expression data analysis. Genome Biol. 19, 15 (2018).

135. Website. 10.2307/2333011 doi:10.2307/2333011.

136. Missarova, A. et al. geneBasis: an iterative approach for unsupervised selection of targeted gene panels from scRNA-seq. Genome Biol. 22, 1–22 (2021).

137. Stringer, C., Wang, T., Michaelos, M. & Pachitariu, M. Cellpose: a generalist algorithm for cellular segmentation. Nat. Methods 18, 100–106 (2021).

138. Gataric, M., et al. PoSTcode: Probabilistic image-based spatial transcriptomics decoder. bioRxiv 2021.10.12.464086 (2021) doi:10.1101/2021.10.12.464086.

139. GitHub - Teichlab/iss_patcher: Approximate missing features from higher dimensionality data neighbours. GitHub https://github.com/Teichlab/iss_patcher.

140. GitHub - Teichlab/visium_stitcher: Stitch multiple Visium slides together. GitHub https://github.com/Teichlab/visium_stitcher.

