## Supplementary figures and images for "Spatiotemporal map of the developing human reproductive tract at single-cell resolution"

### Extended Figure 1

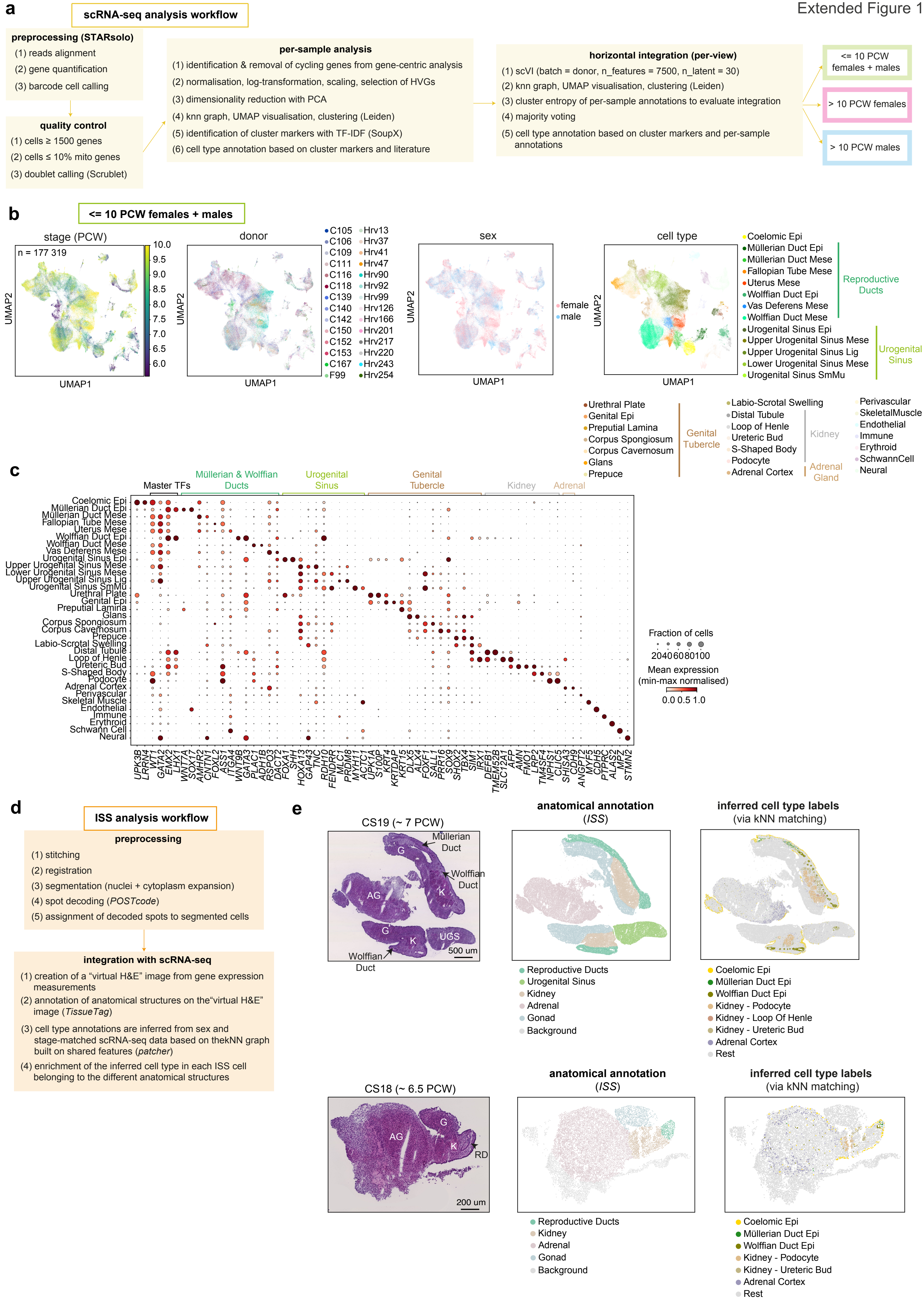

### Extended Figure 2

#### Extended Figure 2

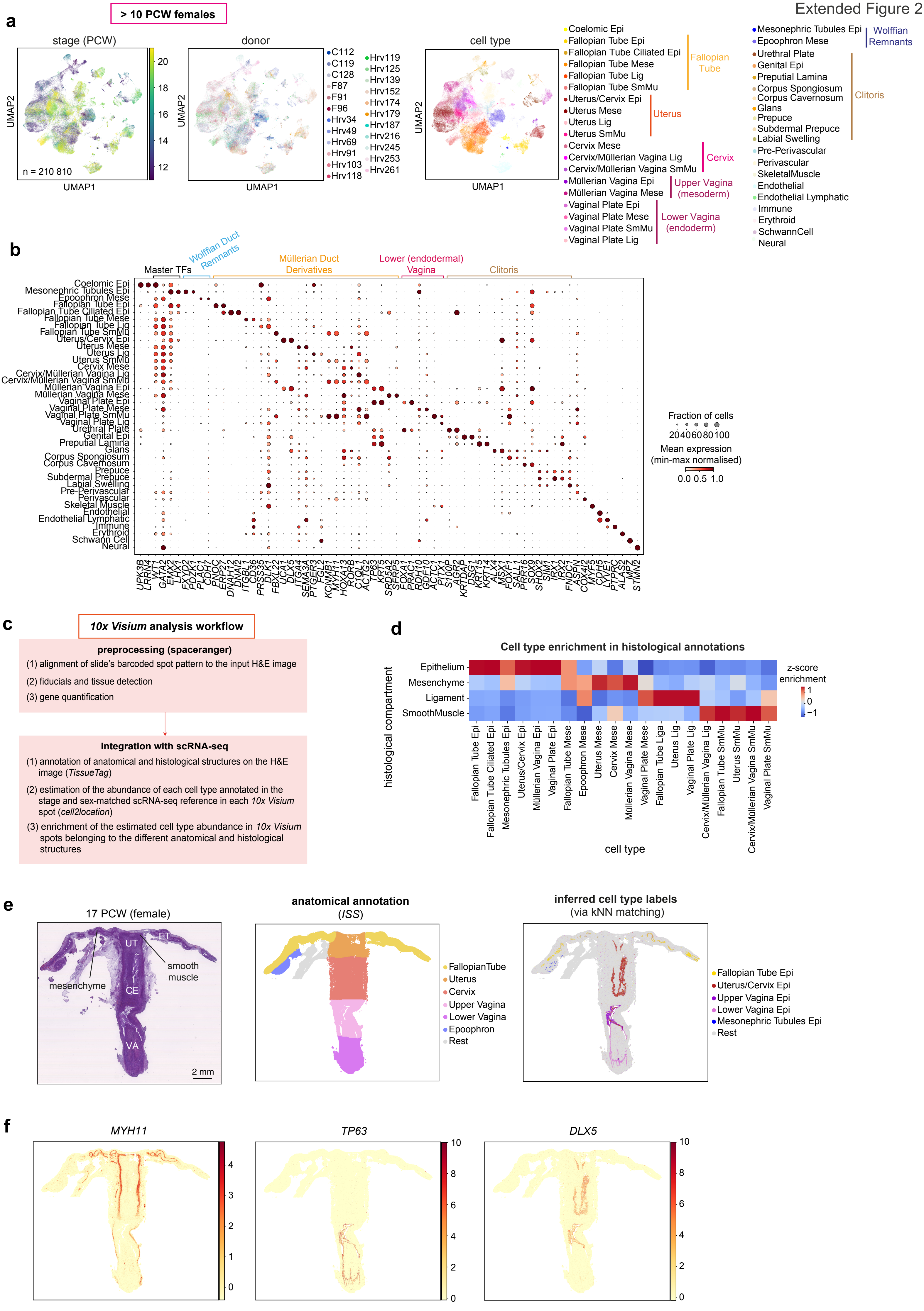

### Extended Figure 3

Extended Figure 3

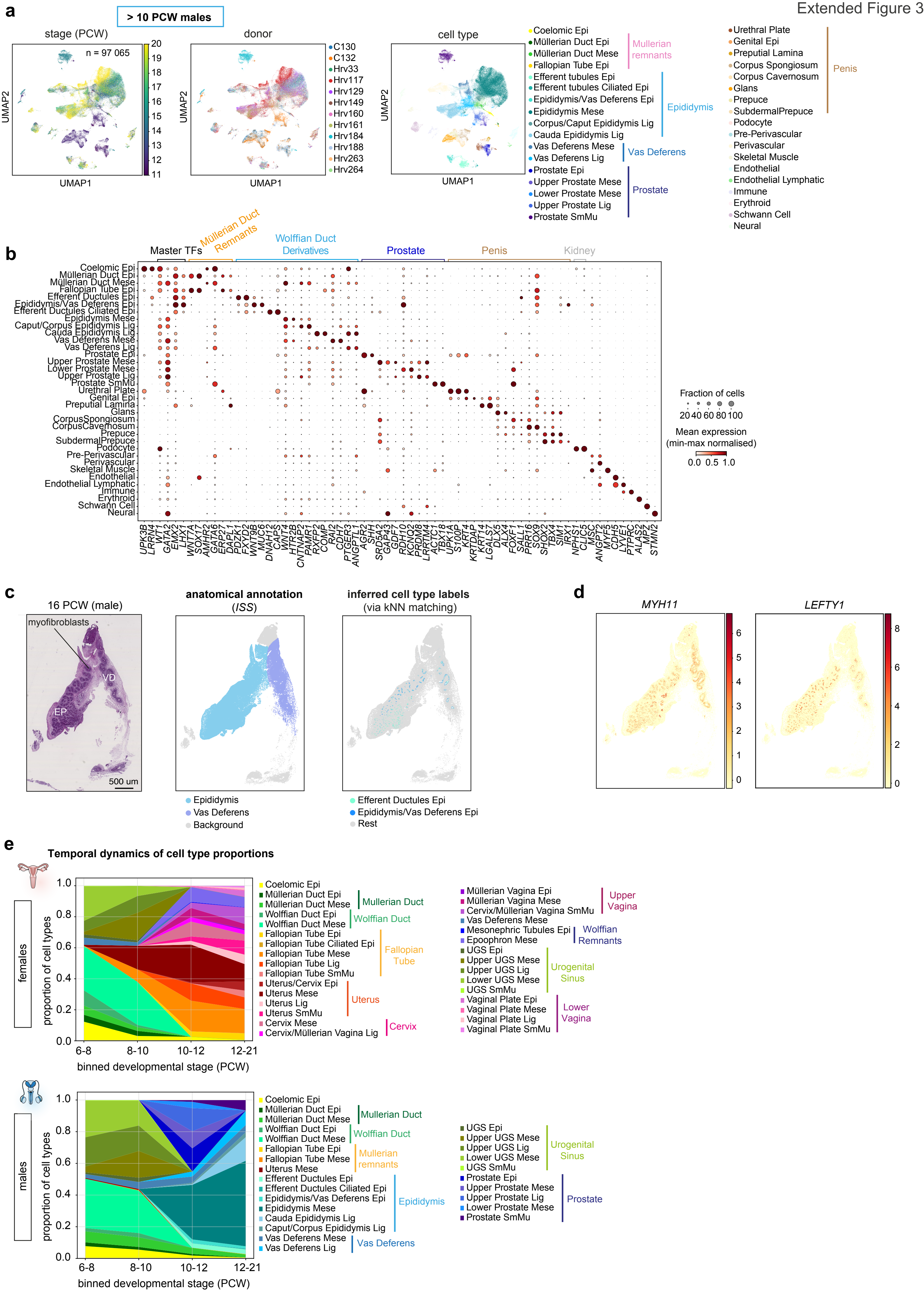

### Extended Figure 4

Extended Figure 4

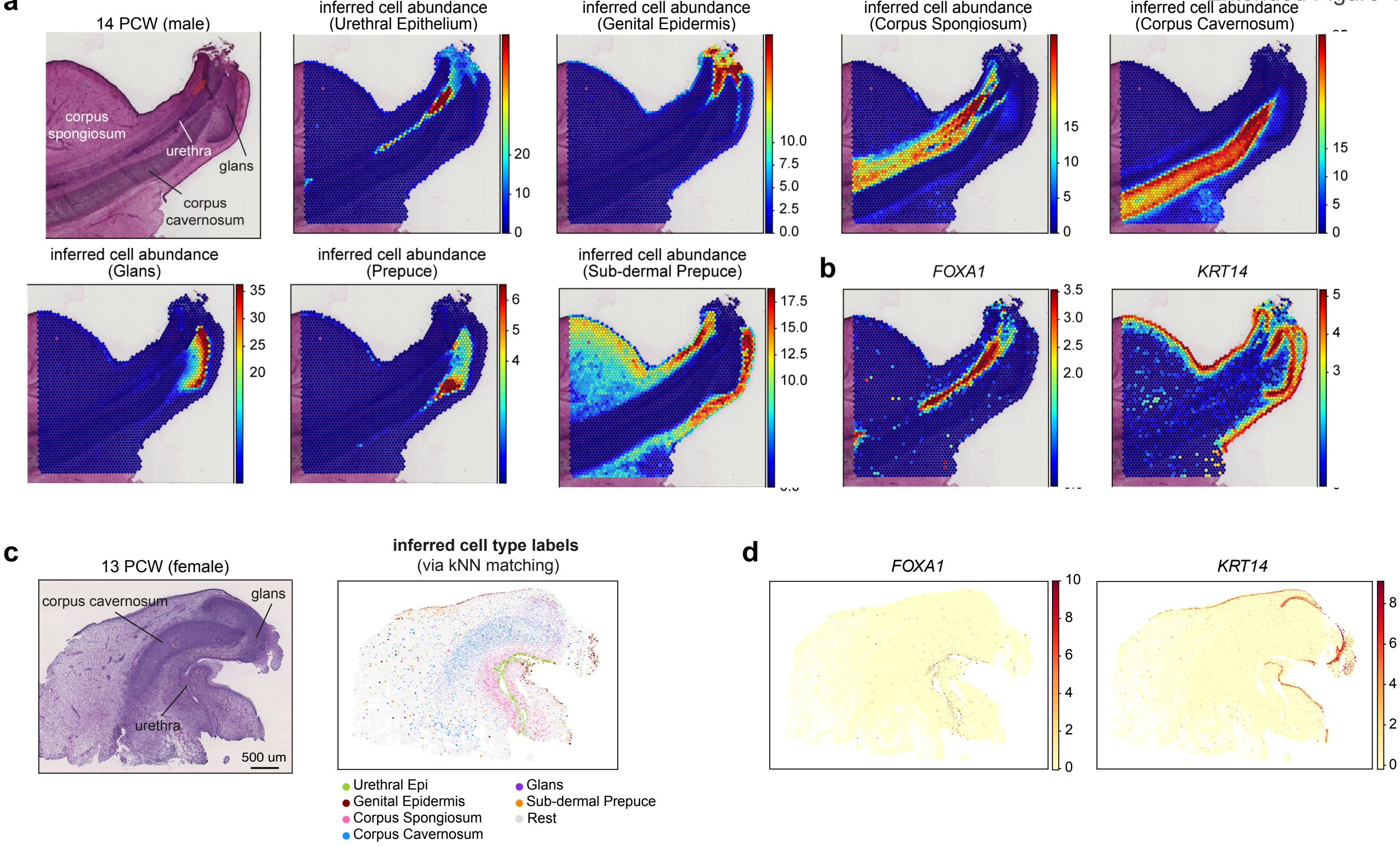

### Extended Figure 5

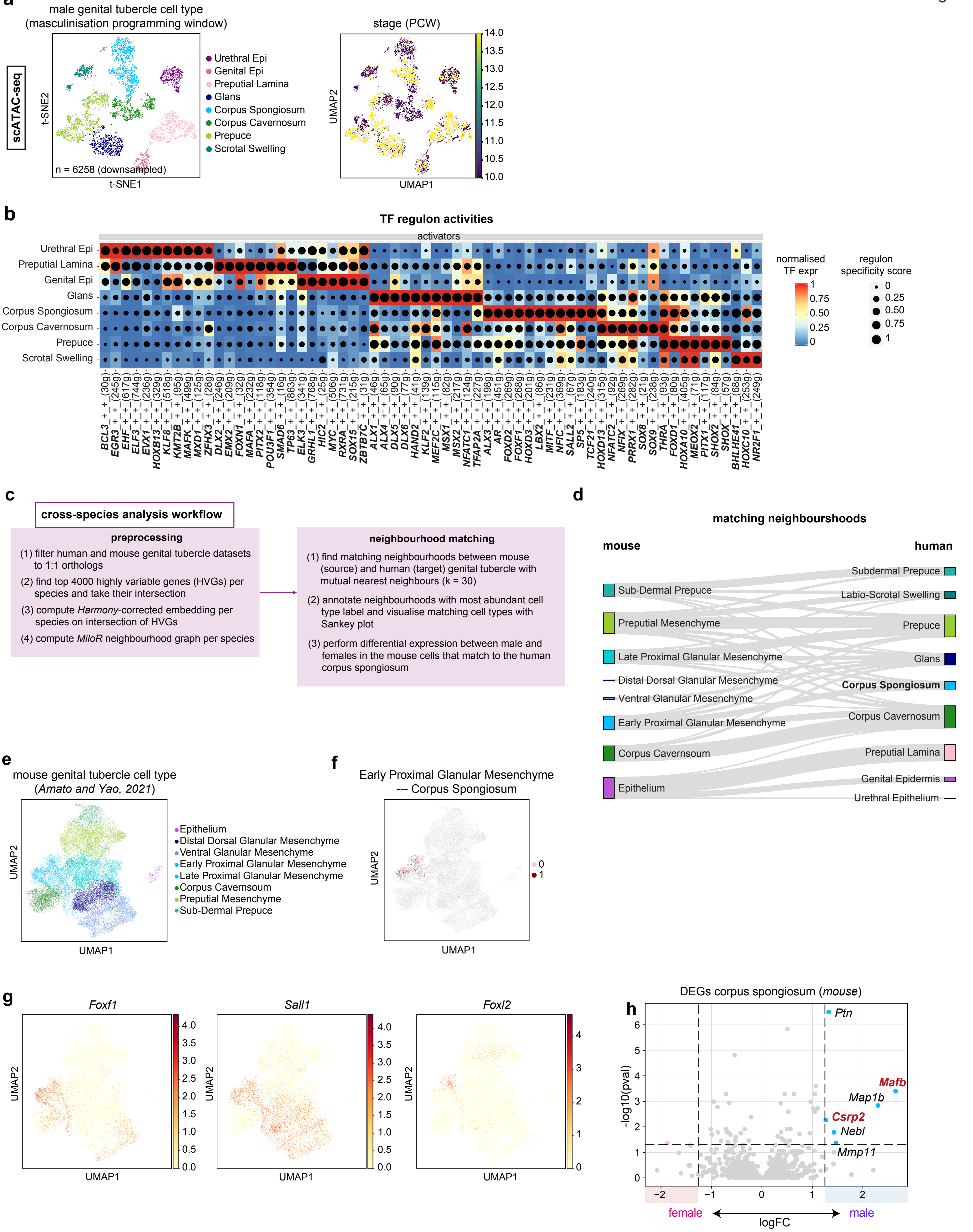

### Extended Figure 6

Extended Figure 6

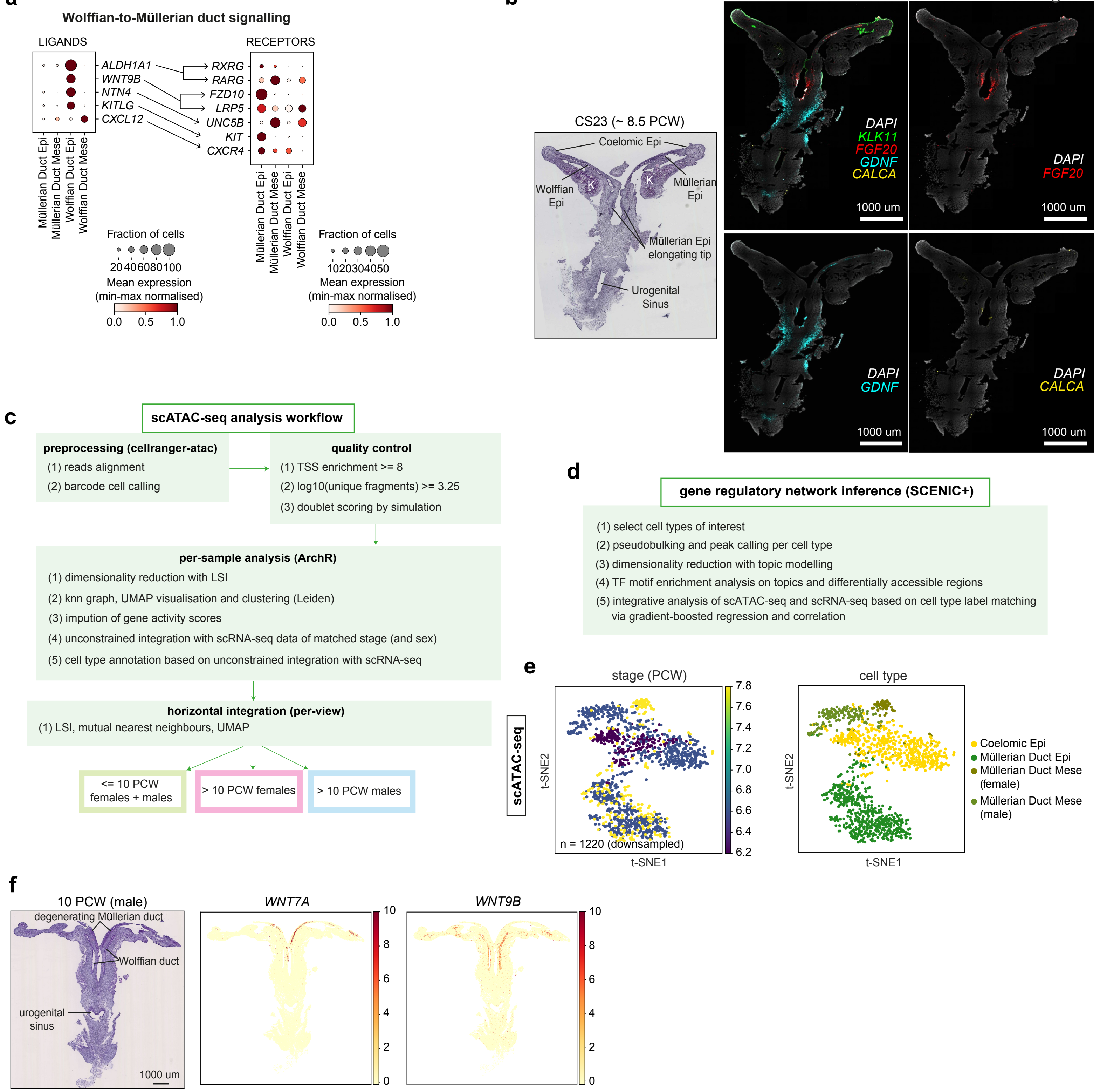

### Extended Figure 7

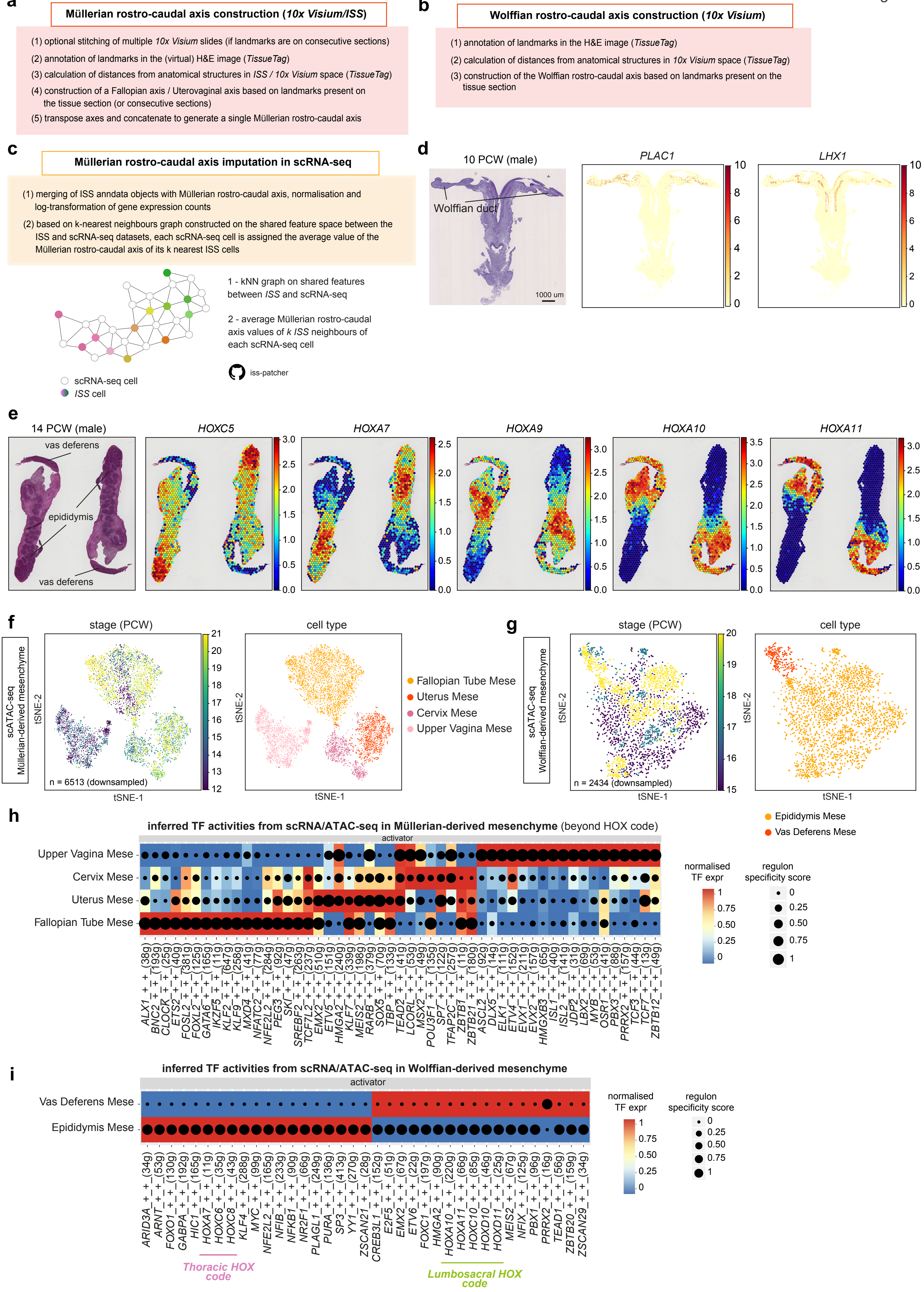

### Extended Figure 8

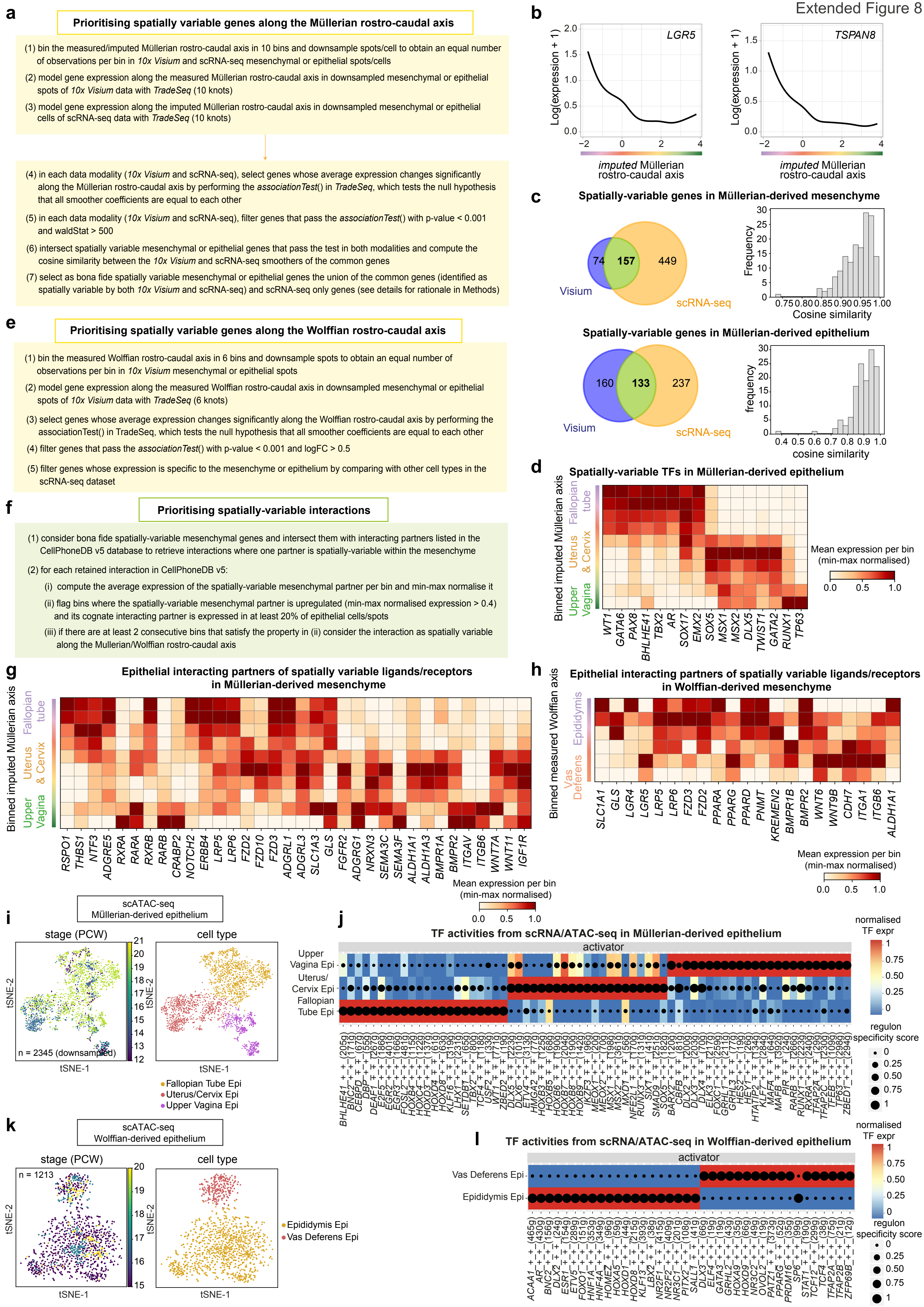
