## Extended Figure 9 for "Spatiotemporal map of the developing human reproductive tract at single-cell resolution"

**a**

**Spatially-variable genes along the Fallopian tube axis**

(1) bin the measured Fallopian tube axis and downsample to obtain an equal number of observations per bin in Visium (only epithelial spots)

(2) model gene expression along the measured Fallopian tube axis in downsampled epithelial spots of Visium data with TradeSeq (5 knots)

(3) select genes whose average expression changes significantly along the Fallopian tube axis by performing the associationTest() in TradeSeq, which tests the null hypothesis that all smoother coefficients are equal to each other

(4) filter genes that pass the associationTest() with p-value < 0.001 and logFC > 0.5

(5) filter genes whose expression is specific to the secretory epithelium by comparing with other cell types in the Fallopian tube using the scRNA-seq dataset

**comparison with adult Fallopian tube**

(6) select epithelial spots in each capture area of the adult Fallopian tube

(7) compute an average score of the increasing and decreasing signature of fetal Fallopian tube genes from fimbria to isthmus in each capture area (using scanpy.tl.score_genes() function)

(7) perform Jonckheere's trend test to check if the increasing and decreasing pattern observed in fetal development is preserved in the adult Fallopian tube

**b**

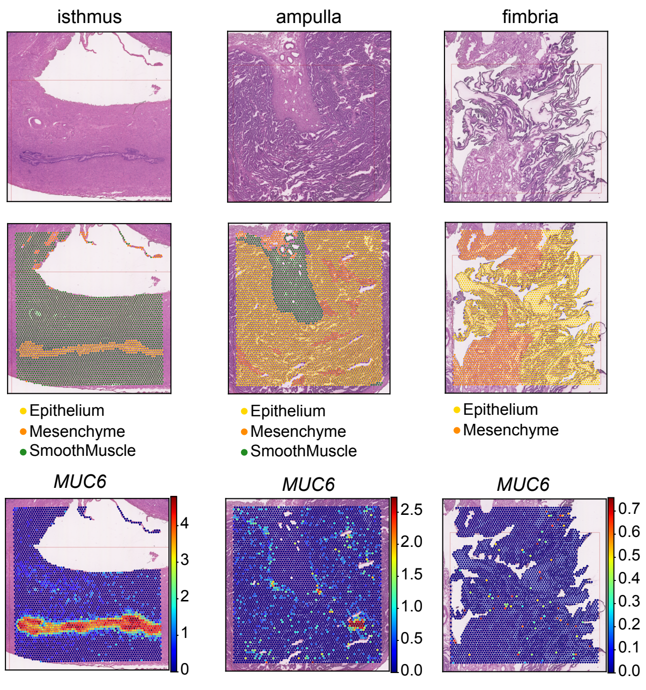


**c**

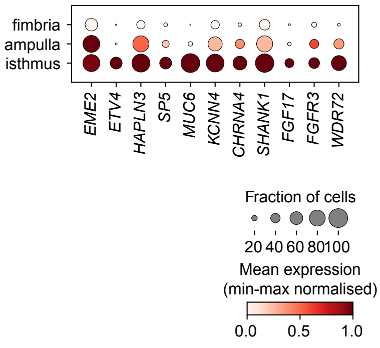


**d**

**Spatially-variable genes along the Epididymal axis**

(1) bin the measured Epididymal axis and downsample to obtain an equal number of observations per bin in Visium (only epithelial spots)

(2) model gene expression along the measured Epididymal axis in downsampled epithelial spots of Visium data with TradeSeq (5 knots)

(3) select genes whose average expression changes significantly along the Epididymal axis by performing the associationTest() in TradeSeq, which tests the null hypothesis that all smoother coefficients are equal to each other

(4) filter genes that pass the associationTest() with p-value < 0.001 and logFC > 0.5

(5) filter genes whose expression is specific to the secretory epithelium by comparing with other cell types in the epididymis using the scRNA-seq dataset. This is particularly important to remove genes of the ciliated epithelium, which is specifically enriched in the rostral portion of the axis (see panel **e**).

**e**

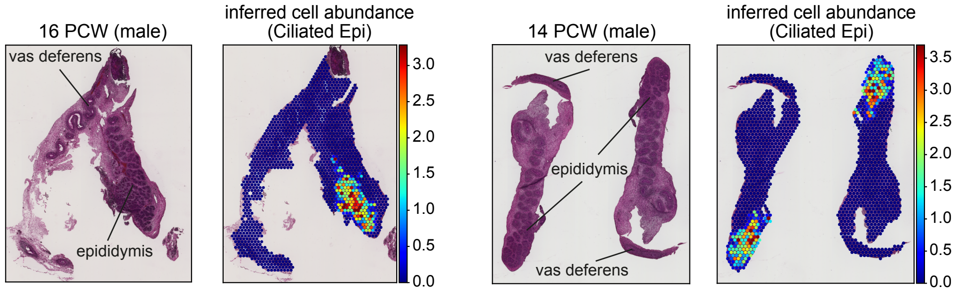
